# Small molecule intervention of actin-binding protein profilin1 reduces tumor angiogenesis in renal cell carcinoma

**DOI:** 10.1101/2025.07.18.665115

**Authors:** David Gau, Pooja Chawla, Kexin Xu, Niharika Welling, Jacob Antonello, Caroline Consoli, Monica Wei, Sara Zdancewicz, Michael DaSilva, Chris Varghese, Paul Francoeur, Haibin Shi, Xucai Chen, Anurag Paranjape, Flordeliza Villanueva, Thomas E. Smithgall, Andrew VanDemark, David R. Koes, Donna M. Huryn, Partha Roy

## Abstract

Angiogenesis plays a key role in the development and progression of renal cell carcinoma (RCC). Actin-binding protein profilin-1 (Pfn1) is overexpressed in clear cell RCC predominantly in tumor-associated vascular endothelial cells (ECs). We previously demonstrated that that EC-selective (over)expression of Pfn1 accelerates RCC progression, and conversely, genetic loss of EC-Pfn1 dramatically inhibits tumor angiogenesis impeding tumor initiation and/or progression in RCC, suggesting that Pfn1 could be an actionable therapeutic target in RCC. In this study, we demonstrate that 4,4’-((4-bromophenyl)methylene)bis(3,5-dimethyl-1*H*-pyrazole), a small molecule that we had previously identified as an inhibitor of Pfn1-actin interaction, directly binds to Pfn1 and attenuates tumor angiogenesis when directly administered into subcutaneous RCC tumors. Next, we undertook a chemical optimization approach to design and synthesize 4,4’-((4-(trifluoromethyl)phenyl)methylene)bis(3,5-dimethyl-1*H*-pyrazole), a structural analog of our originally identified inhibitor, that exhibits improved anti-angiogenic efficacy *in vitro* and *in vivo*. Finally, we demonstrate that Pfn1 inhibitor is amenable to lipid microbubble encapsulation and release in the tumor microenvironment (TME) by ultrasound-mediated disruption of circulating microbubbles to achieve anti-angiogenic and anti-tumor benefit. In summary, our findings suggest that tumor-localized release of Pfn1 inhibitor could be a potential therapeutic strategy in RCC.

## INTRODUCTION

Renal Cell Carcinoma (RCC) is the most common form of kidney cancer and among the different histological subtypes of RCC, clear cell RCC (ccRCC) occurs in >75% of RCC patients. Although surgical and ablative procedures successfully cure early-stage ccRCC, approximately 30% of ccRCC patients present with metastasis at the time of diagnosis (1). Another one-third of patients, following initial treatment, develop either local recurrence and/or distant metastases with a 10% five-year survival expectation. ccRCC is characterized by near-universal loss of the entire or most of the 3p chromosome, resulting in genetic loss-of function (LOF) of the ubiquitin ligase *Von Hippel Lindau* (VHL) in >90% cases and other chromatin-remodeling and DNA-repair genes (e.g. PBRM1, BAP1, SETD2) at various frequencies in progressor tumors. A major consequence of LOF of VHL is hypoxia-inducible factor (HIF)-mediated transcriptional upregulation of various pro-angiogenic growth factors (VEGF, PDGF) leading to a highly vascularized tumor microenvironment (TME) and tumor progression (2). Although anti-angiogenic therapies (AATs) combined with immunotherapy are front-line treatments for ccRCC (3), resistance to AATs remains a persistent clinical problem for ccRCC patients (4–6).

Endothelial cell (EC) migration and proliferation are essential processes of angiogenesis that are highly dependent on the action of the actin cytoskeleton (7). Actin monomer-binding protein Profilin1 (Pfn1 – the most abundant and ubiquitously expressed protein within the Pfn family) plays an important role in facilitating actin polymerization in cells. Depletion of Pfn1 generally results in reduced cellular levels of polymerized actin (F-actin), causing defects in proliferation and migration in a variety of cell types including ECs (8–12). We previously reported that conditional knockout (KO) of the *Pfn1* gene in vascular ECs causes major defects in developmental angiogenesis (9). Angiogenic stimulation increases Pfn1 phosphorylation in ECs in a site-specific manner (Y129), and genetic abrogation of Y129 phosphorylation of Pfn1 leads to reduced angiogenesis and tumor progression in a glioblastoma (GBM) setting (13, 14). These findings underscore endothelial Pfn1’s important role in both physiological and pathological angiogenesis.

Pfn1 expression is transcriptionally elevated in both clear cell and papillary RCC which together account for >90% of RCC cases (15, 16). For ccRCC, Pfn1 is one of the top-12 differentially (over)-expressed proteins in metastatic *vs*. primary RCC(17), and Pfn1 overexpression is linked to adverse clinical outcomes (shorter overall as well as progression-free survival) in human ccRCC (16–20). In ccRCC, we previously reported that Pfn1 expression is more commonly dysregulated in stromal cells rather than in tumor cells, most notably in tumor-associated vascular ECs (16). Supporting these clinical correlative findings, we further showed that endothelial (over)expression of Pfn1 accelerates orthotopic tumor progression of RENCA (a commonly used murine VHL+ RCC cell line) cells in syngeneic mouse models. Conversely, triggering EC-specific loss of Pfn1 dramatically abrogates tumor formation and/or progression of RENCA tumors, and importantly, Pfn1-dependency for tumorigenicity of RENCA cells was maintained even in the absence of VHL. As expected, RCC tumors with genetic LOF of EC Pfn1 exhibited a dramatic reduction in tumor angiogenesis (15). Collectively, these findings suggest that Pfn1 represents a promising therapeutic target for diseases driven by excessive neovascularization such as RCC.

In our previous work, we discovered two structurally distinct non-cytotoxic small molecule compounds, namely, 8-(3-hydroxyphenyl)-10-phenyl-2,4,5,6,7,11,12-heptaazatricyclo[7.4.0.0^3^,7] trideca-1(13),3,5,9,11-pentaen-13-o (denoted as compound C2) and 4,4’-((4-bromophenyl)methylene)bis(3,5-dimethyl-1*H*-pyrazole) (denoted as compound C74) that reversed Pfn1’s effect on actin polymerization in biochemical assays. Treatment with these compounds reduced Pfn1:actin interaction in cells, inhibited cell migration and proliferation in a dose-dependent manner, and diminished abnormal ocular neovascularization *in vivo* (9, 16, 21, 22). Tumor angiogenesis is a complex process that involves the actions of multiple different cell types in the tumor microenvironment (TME) including ECs, tumor cells, and other types of stromal cells such as cancer-associated fibroblasts and various types of tumor-infiltrated immune cells. Whether tumor angiogenesis is also susceptible to small molecule-mediated intervention against Pfn1-actin interaction in the TME has remained unexplored. In the present study, we established the utility of C74 (biologically more potent than C2 in inhibiting cell migration and proliferation (16)) as an anti-angiogenic agent in RCC tumors, and report the design and synthesis of a new structural analog of C74 with improved anti-angiogenic efficacy *in vitro* and *in vivo*. Furthermore, we demonstrate the feasibility of tumor-localized delivery of Pfn1 inhibitor by an ultrasound-guided method to achieve reduction in tumor angiogenesis and tumor growth.

## MATERIALS AND METHODS

### Cell Culture and Reagents

RENCA renal cell carcinoma (RCC) cells (ATCC, CRL-2947) were cultured in RPMI supplemented with 10% fetal bovine serum (FBS) and 1% penicillin-streptomycin. Human microvascular ECs (HmVECs, ATCC, CRL-3243) were cultured in MCDB-131 media with growth supplements as described previously (21). Cell lines were tested for mycoplasma contamination.

### Small molecule compounds

C74 was either purchased from commercial sources (MolPort-000-793-534) or chemically synthesized. Other analogs were synthesized as described in the ***Supplementary Information* (SI)** section. All tested compounds were determined to be of at least 90% chemical purity. The details of chemical synthesis of various C74 analogs and structural characterization are provided in the ***SI*** section.

### Molecular docking

Molecular docking was performed using GNINA (23) which performs Monte Carlo search over possible ligand conformations and poses and ranks the resulting poses using a convolutional neural network (24). Default settings were used and the protein receptor structure was kept rigid. The protein structure used for docking includes a cryptic pocket at the Pfn1:actin interface that was identified through molecular dynamics simulations, as previously described (22).

### Pfn1 purification and Differential Scanning Fluorometry (DSF) assay

Recombinant wild-type (WT Pfn1 was expressed in BL21(DE3) Codon + RPIL *E. Coli* as an N-terminal fusion with a purification tag containing both αν 8X-His and the fluorescent protein, mRuby2. After lysis, the fusion protein was purified using IMAC (immobilized metal affinity chromatography) and the His_8_-mRuby2 tag liberated from the fusion protein by overnight digestion with TEV (Tobacco Etch Virus) protease. The purification tag and TEV protease were then separated from Pfn1 using a second round of IMAC. The resulting Pfn1 is then subjected to ion exchange and size exclusion chromatography. The resulting purity is >99% as judged by SDS-PAGE. For the DSF assays, Pfn1 was used at a concentration of 0.3 mg/ml in a buffer containing 50mM HEPES pH7.5, 50mM NaCl, 1 mM β-mercaptoethanol, and 5X SYPRO Orange. Fluorescence of SYPRO Orange was measured and plotted as a function of temperature to quantify protein unfolding. The T_m_ was defined as the temperature with the maximum rate of change in fluorescence.

### Surface Plasmon Resonance

SPR analysis was conducted using a Reichert 4SPR instrument (Reichert Technologies) with 10 mM HEPES, pH 7.4, 150 mM NaCl, 0.05% Tween 20, and 3 mM EDTA as the running buffer. An anti-GST antibody (Sigma Aldrich, G7781) was immobilized first on carboxymethyl dextran hydrogel biosensor chips (Reichert) using standard amine coupling chemistry with 1-ethyl-3-(-3-dimethylaminopropyl) carbodiimide hydrochloride (EDC) and N-hydroxysuccinimide (NHS). Purified recombinant GST-Pfn1 proteins or GST alone as a negative control were then captured by the immobilized antibodies on separate channels on the chip. Analytes (purified rabbit muscle actin (source: Cytoskeleton Inc., Denver, CO), C74 or its analog) were then injected over a range of concentrations at 50 µL/min with a 60 s association phase and 120 s dissociation phase. The presence or absence of Mg^2+^ does not impact the K_d_ value of the Pfn1-actin interaction; however, the critical concentration of actin polymerization dramatically rises from its usual sub-μM (in the presence of 2 mM MgCl_2_) ranges to values in the tens of μM in the absence of Mg^2+^ (25). Therefore, to ensure that the injected actin remains in monomeric form throughout the actin concentration range (0.1-3 μM) of the SPR studies, actin was pre-dialyzed against the SPR running buffer that is free of MgCl_2_. The chip surface was regenerated with 1 mM NaOH after each analyte injection. Each analyte concentration was measured in duplicate (actin) or triplicate (C74), and the resulting sensorgrams were corrected for buffer effects. Kinetic and binding constants were calculated by fitting the sensorgrams with a 1:1 Langmuir binding model using the TraceDrawer software (Reichert).

### In vitro anti-proliferation and anti-angiogenesis screening

For initial screening of compounds in a cell proliferation assay, HmVECs were seeded in the wells of a 24-well plate and treated with various compounds at a 20 μM concentration or equivalent concentration of DMSO (solvent control), and cell count was measured well-by-well by imaging and extrapolating the overall number of cells/well using a Digital Cell Imager (Millipore). For testing of anti-angiogenic capability of small molecules, HmVECs were plated on growth factor-reduced matrigel (R&D Systems, 3433-005-01) and treated with various compounds at a 20 μM concentration (unless indicated otherwise) or equivalent concentration of DMSO as described previously (21). Endothelial cord formation was imaged by either phase-contrast or fluorescence (where cells were pre-labeled with Calcein-AM green (Invitrogen, Carlsbad, CA)) imaging and quantified after 18 hours, with the length of branch segments used as a measure of angiogenic activity.

### Lipid microbubble encapsulation of C74 and UP6 (C74 analog)

Lipid microbubbles were prepared with modifications to a previously described method (26). DSPC (1,2-distearoyl-sn-glycero-3-phosphocholine; 850365C, Avanti Polar Lipids, Alabaster, AL, USA), DSPE-mPEG 2000 (1,2-distearoyl-sn-glycero-3-phosphoethanolamine-N-[methoxy(poly-ethylene glycol)-2000] ammonium salt; 880120C, Avanti Polar Lipids), and poly-oxyethylene (40) stearate (P3440, Millipore Sigma) dissolved in chloroform were mixed with C74 or UP6 (dissolved in ethanol) in a 2:1:1:0.5 (w/w/w) ratio and dried under argon at room temperature for 25 to 30 minutes, followed by overnight vacuum drying. The dried lipid film was hydrated with saline, and the resulting dispersion was sonicated with a Misonix XL2020 Sonicator Ultrasonic Processor XL (Bioventus, Farmingdale, NY, USA) at power level 5.25 in the presence of perfluorobutane (PFB) gas (FluoroMed, Round Rock, TX, USA) for 1 minute 15 seconds to form microbubbles. These were washed twice in a saline bag, incubating at room temperature for 60 minutes after each wash, then resuspended in saline. The final formulation was aliquoted into vials with PFB-filled headspace and stored at 4 °C until use. This method consistently produced microbubbles with an average concentration of ∼0.5 × 10⁹/mL and a mean diameter of 3 µm, as measured using a Coulter counter (Multisizer 4e, Beckman Coulter, Indianapolis, IN, USA) with a 30 μm aperture tube. Quantitative mass-spectrometry (MS) of compound (C74 or UP6)-loaded microbubble solution (microbubbles were ruptured by vortexing to release compounds prior to performing MS) against known compound standard amount were performed at the MS facility of Scripps Research to estimate the compound concentration in the microbubble solution (= standard concentration x area under microbubble solution MS curve / area under standard MS curve) (*section E* of **Supplementary Information**).

For *in vitro* testing of microbubble-encapsulated C74, HmVECs were seeded for cord formation as described above and allowed to grow for 24 h. Microbubbles loaded as indicated were added to the culture media to achieve a final ratio of 10 microbubbles per cell. The wells were covered with cell media, sealed with polycarbonate membrane and inverted. To disrupt microbubbles and release C74, Ultrasound was delivered with a single-element immersion transducer (A302S, 25.4 mm in diameter, Olympus NDT, Center Valley, PA), driven by an arbitrary function generator (AFG3252, Tektronix, Beavertona, OR) connected to a gated radio frequency power amplifier (250A250AM8, Amplifier Research, Souderton, PA). The ultrasound field was calibrated with a 200-µm capsule hydrophone (HGL-0200, Onda Corp, Sunnyvale, CA). Ultrasound pulses were delivered at 1 MHz, 0.50 MPa peak negative pressure (spatial peak temporal peak), 10 µs pulse duration, and pulse interval 1 ms (duty cycle 1%), for a total of 10 s. The media was removed after ultrasound-targeted microbubble destruction (referred to as UTMD heron), and cells were replenished with 10% DMEM and incubated for 48 hr for cell viability assay.

### In vivo studies

#### i. Matrigel plug angiogenesis assay

A single dose of 0.5 ml of growth factor reduced matrigel plugs supplemented with bFGF (500 ng) and either DMSO or compound (200 μM) was subcutaneously implanted bilaterally into the flanks of C57/B6 mice (5-6-week-old male mice). After 10 days, the plugs were excised, and infiltrating ECs into the plugs were identified by CD31 immunostaining and quantified to assess the extent of vascularization.

##### RCC tumor model studies

All animal experiments were conducted in compliance with an approved IACUC protocol, according to University of Pittsburgh Division of Laboratory Animal Resources guidelines. As previously described (16), 1 million RENCA cells were injected in a 1:1 PBS/matrigel mix subcutaneously into the flanks of 6–7-week-old male immunocompetent Balb/C mice to establish RCC tumors. For direct intratumoral injection, C74 (16 mg/kg) or equivalent amount of DMSO dissolved in saline was injected at the tumor cell inoculation site over a course of 19 days starting on day 1 before sacrificing the animals. For studies involving microbubble-based delivery of C74 or UP6, subcutaneous RENCA tumors were established as described above. One hundred μl suspension of microbubbles in PBS was retro-orbitally injected into animals. Ultrasound pulses were delivered for 25 minutes with a clinical ultrasound imaging system (S3 probe, Phillips Sonos 7500). The ultrasound probe was placed on the tumor, and the ultrasound was operated in ultra-harmonic mode (center frequency of 1.3 MHz) with an on-screen mechanical index of 1.6. The system was time-triggered, with 4 frames per burst and a burst interval of 2 s to allow reperfusion of the tumor with microbubbles in between bursts. The treatment was monitored by imaging the tumor using a 15L8 transducer probe on a Sequoia 512 ultrasound imaging system (Siemens Ultrasound, Issaquah, WA, USA) operated in Contrast Pulse Sequencing (CPS) mode (mechanical index = 0.20 and frame rate of 5 Hz) to confirm MB destruction by therapy pulses and subsequent reperfusion of MBs in the tumor. For tumor growth inhibition studies (N = 8-9 mice per group), treatment was initiated after 5 days post inoculation and animals received a total of three UTMD treatments at two to three-day intervals. Mice were euthanized 2 days after the last treatment and tumors were harvested.

### Immunohistochemistry

Tissue sections were deparaffinized and rehydrated before blocking and incubating overnight with an anti-CD31 antibody (77699, Cell Signaling Technology, 1:100). For immunodetection of the primary antibody, a secondary biotin-labeled anti-rabbit antibody (ab97049, Abcam) and streptavidin-peroxidase conjugates (11089153001, Roche) were used. The staining was detected with the Pierce DAB substrate kit (34002, Thermo Fisher). For some experiments, the slides were counterstained with hematoxylin before dehydration and mounting.

### Statistics

Statistical tests were performed with either one-way ANOVA followed by Tukey’s post hoc test or nonparametric Mann–Whitney test when appropriate unless specified otherwise. Differences exhibiting p <0.05 were considered as statistically significant.

## RESULTS

### Demonstration of C74 as an inhibitor of tumor angiogenesis

In a RENCA-based RCC mouse model, we previously showed that direct intratumoral administration of C74 (chemical structure shown in **Figure 1A**) leads to reduced tumor growth in a subcutaneous implant setting (16). However, since our *in vitro* studies also showed that C74 treatment reduces RENCA cell proliferation, we could not discern the relative contributions of tumor-intrinsic vs -extrinsic effects of C74 underlying its anti-tumor action. Therefore, we performed an end-point assessment of tumor angiogenesis by performing immunohistochemistry (IHC) of tumor sections for CD31 (a marker of vascular ECs) and quantifying the relative CD31-positive areas in DMSO (vehicle control)- vs. C74-treated RENCA tumors collected from our previous *in vivo* studies. As per these analyses, C74 treatment reduced the CD31-positive area by ∼30% in a statistically significant manner (**Figure 1B-C**). These data suggest that C74 is capable of inhibiting tumor angiogenesis.

**Figure 1.**
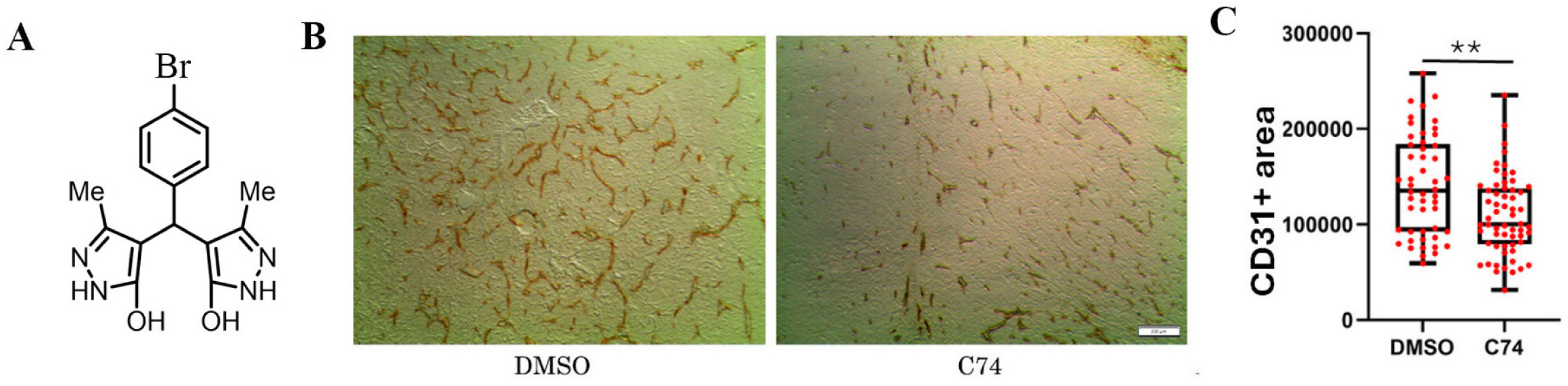
Demonstration of C74’s ability to diminish tumor angiogenesis. **A)** Chemical structure of C74. **(B-C)** Representative images (*panel B*; 4× magnification) and quantification (*panel C*; based on 10× field images) of CD31^+^ cells in subcutaneously established RENCA tumors subjected to daily direct intra-tumoral injection for 19 days with either C74 or equivalent amount of DMSO control. in BALB/c mice (two-tailed Student’s T-test, **, p < 0.01; n = 5 mice/group; scale bar, 200 μm).

### Biochemical characterization of C74

While Pfn1’s action enhances actin polymerization in cells, in biochemical assays without the presence of any other actin-assembly factors, Pfn1 inhibits actin polymerization through preventing actin incorporation onto the pointed ends of growing actin filaments as well as by inhibiting spontaneous actin nucleation. Our discovery of C74 as an inhibitor of the Pfn1-actin interaction was based on the results of a pyrene-actin polymerization screening assay where C74 reversed recombinant GST-Pfn1’s inhibitory effect on actin polymerization but had no discernible effect on actin polymerization on its own (16). In the same study, we further demonstrated that C74 treatment reduces proximity-ligand interactions of endogenous Pfn1 and actin in cells and induces other phenotypic changes (i.e. reduced cell spreading, migration, and proliferation) that are expected upon inhibition of cellular Pfn1-actin interaction. However, whether C74 directly binds to Pfn1 and with what affinity has remained unexplored. To address this question, we performed SPR studies with C74 and immobilized GST-Pfn1 (GST served as a negative control). GST epitope-tagging enabled us to immobilize all purified proteins with an anti-GST antibody on the SPR chip thereby minimizing the masking of any binding site of Pfn1 for potential C74’s interaction. As an initial test for the utility of the SPR assay to detect known ligand interactions of Pfn1, we confirmed GST-tagged wild-type (WT) Pfn1 interacts with G-actin while GST-tagged H119E-Pfn1 (a point-mutant of Pfn1 that has ∼25-fold reduction in actin-binding vs WT-Pfn1(27, 28)) did not exhibit any detectable interaction with G-actin as expected (**Figures 2A-B, E**). The estimated dissociation constant (K_d_) of GST-Pfn1:actin interaction in the SPR assay was approximately equal to 5.3 μM (**Figure 2E**). This value is very much in agreement with the K_d_ value (= 5 μM) that was previously measured for human Pfn1’s interaction with rabbit muscle actin (as used in our study) by Goldschmidt-Clermont and colleagues by three independent methods (29) as well as the general range of K_d_ values (1-10 μM) reported by other investigators (30–32). We were able to detect C74’s interaction with Pfn1 with an estimated K_d_ value approximately equal to 60 μM; however, H119E substitution did not impair C74’s interaction with Pfn1 (**Figures 2C-E**). This K_d_ value is consistent with the concentration requirement of C74 in the tens of micromolar range to achieve biological effects as we previously demonstrated in angiogenesis assays *in vitro* (21). These results suggest that C74 is a low-affinity binder of Pfn1, further motivating us to identify a more potent analog of C74 by medicinal chemistry approach as described in the following section.

**Figure 2.**
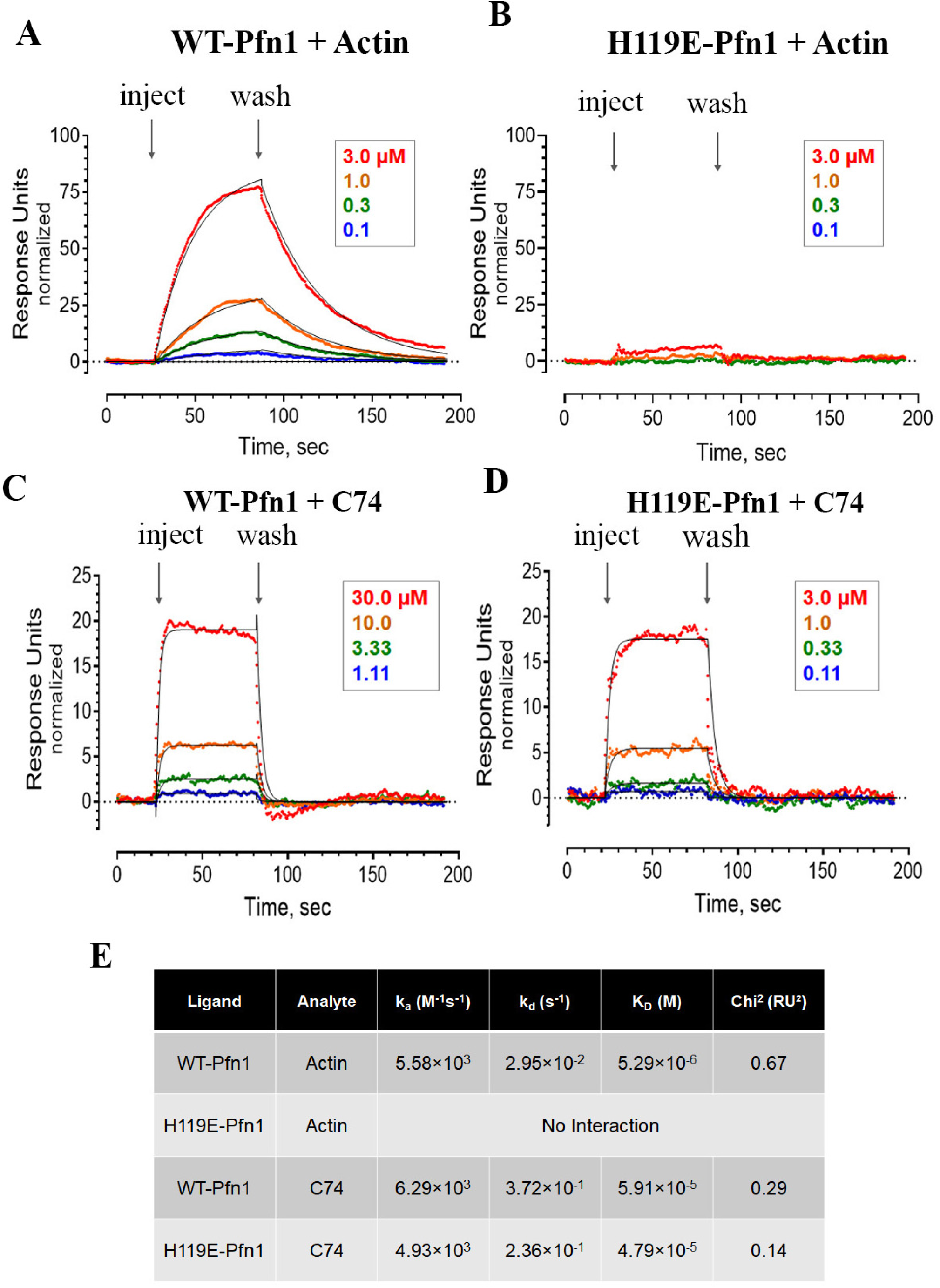
SPR-based demonstration of C74-Pfn1 interaction. **A-E**) Recombinant purified wild-type (WT) and H119E mutant forms of Pfn1 were immobilized on an SPR chip as described under Materials and Methods. Purified Actin (*panels A, B*) or C74 (*panels C, D*) were then injected over the range of concentrations shown either in duplicate (Actin) or triplicate (C74) followed by a dissociation phase (*arrows*). The resulting sensorgrams are shown with the data points in color and the fitted curves in black. Response units are normalized to the amount of Pfn target protein immobilized on each channel. The reference-corrected sensorgrams were fit to a 1:1 Langmuir binding model and the resulting kinetic constants are shown in the Table (*panel E*). K_D_ values were calculated from the relationship K_D_ = k_d_/k_a_.

### Optimization of C74 identifies a new analog with improved anti-angiogenic activity *in vitro* and *in vivo*

In a previous study, we established that the bromine (Br) substituent in C74 could be effectively replaced with either F or CH_3_, and these analogs exhibited anti-angiogenic activity at 10 μM, an improvement over C74 (21). Based on these promising results, we sought to identify additional analogs with improved potency, efficacy and drug-like properties. Based on our docking studies of C74 (**Figure 3A**), we focused on the aryl group as its binding pocket was not fully occupied, and our previous studies supported that modification at this position was not only tolerated but led to improvements in potency, possibly through interactions with R74 and H119. In addition, the pyrazole was predicted to engage in multiple H-bonding interactions with the target, and therefore, changes to this position were likely to lead to less active compounds.

**Figure 3.**
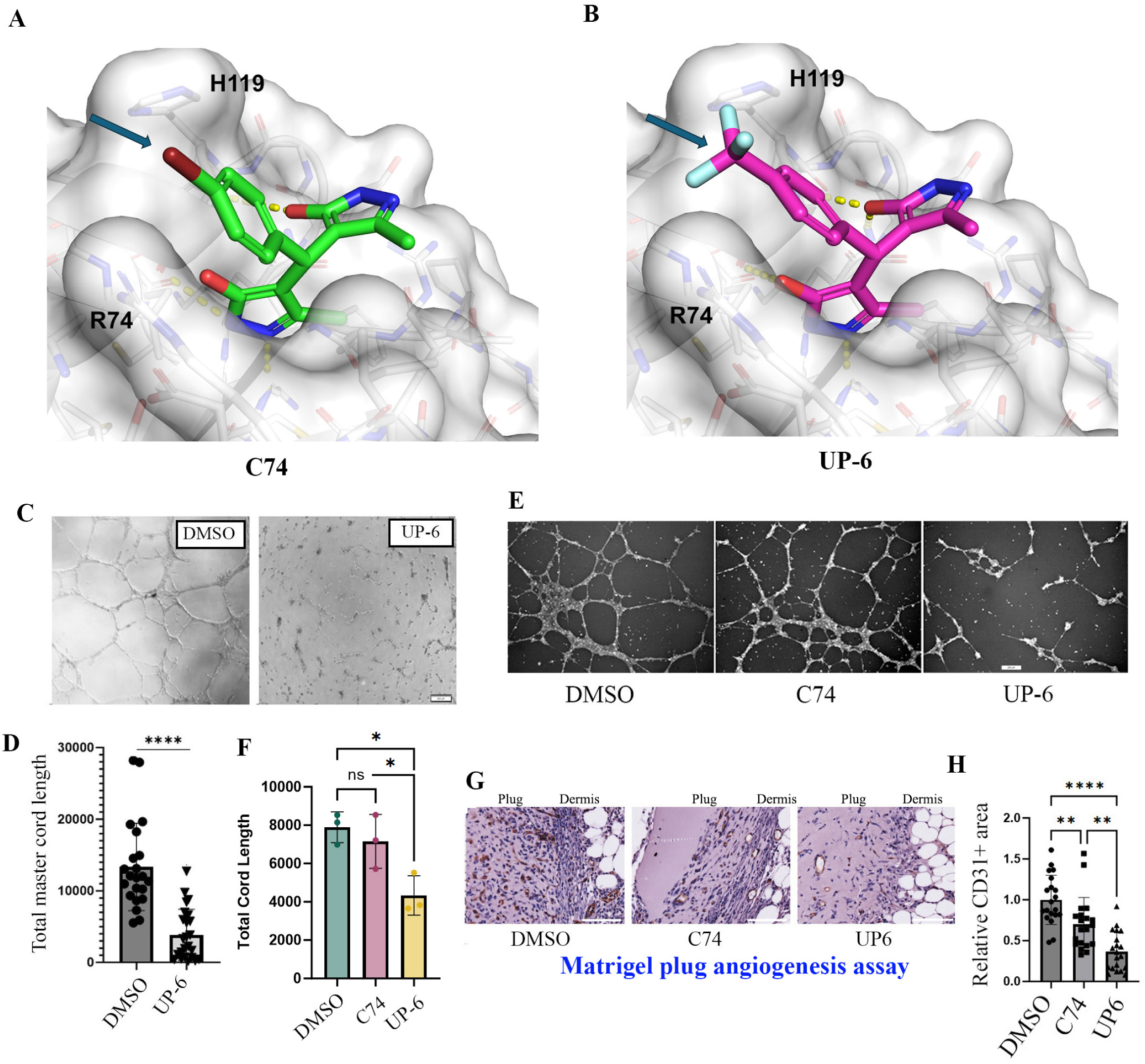
Discovery of a novel analog of C74 with superior anti-angiogenic activity *in vitro* and *in vivo*. **A-B)** Docking models of C74 and UP-6 interaction with Pfn1 - The binding modes of C74 (*panel A*) and UP-6 (*panel B*) as predicted by GNINA. Blue arrow indicates the altered structure on each compound. The unmodified hydroxypyrazoles are predicted to make an identical network of hydrogen bonds in both compounds while the modified aryl ring can potentially interact with R74 and H119. **C-D)** Representative images (*panel C*; 4x magnification) and quantification (*panel D*; based on 4x field images) of cord formation by HmVECs on Matrigel treated with either 20 μM compound UP6 or equivalent DMSO control. Cord length is determined by distance between each cord junction, i.e. cord intersections. Each datapoint represents quantification of cord formation in a single field of observation (two-tailed Student’s T-test; ****, p < 0.0001; n = 3 experiments; scale bar, 200 μm). **E-F)** Representative fluorescence images (*panel E*) and quantification (*panel F*) of cord formation by HmVECs (labeled with cell tracker dye) subjected to 10 μM of either C74 or UP6 vs DMSO (control) (One-way ANOVA with Tukey multiple comparison post-hoc test, * - p<0.05; ns – not significant; scale bar, 200 μm). **G-H**) Representative images of CD31 immunohistochemistry with hematoxylin counterstaining (*panel G*; 4× magnification) and quantification (*panel H*; based on 20X field images) of CD31+ cell infiltration in the subcutaneously implanted Matrigel plugs in Balb/C mice co-injected with a single 200 µM dosage of either C74 or UP-6 or DMSO control. Data are summarized from quantification of 10 bilaterally injected plugs (n=5 mice) per treatment group (One-way ANOVA with Tukey post-hoc test, **, p < 0.01; ****, p < 0.0001; scale bar, 200 μm).

Based on the very limited SAR available (e.g. F and CH_3_ substitution led to greater efficacy than Cl or H) we designed potential new analogs based on: 1) docking studies that suggested the aryl bromine pocket was not fully occupied, 2) exploring whether lipophilicity, polarity or electronegativity contributed to activity, and 3) optimizing the physical chemical properties such as solubility and permeability. A series of potential analogs were docked using the GNINA molecular docking software (23), and we used the docking scores and other factors (e.g. predicted interactions and physical chemical properties) to prioritize compounds for synthesis. Examples are shown in **Table 1** (See **Supplementary Information** *Sections A-D* for synthetic protocols and characterization). The aryl halide was replaced with various pyridine isomers (UP-1, −4, and −5) to evaluate if electronegativity contributed to binding and to potentially enhance solubility; CF_3_ (UP-2, −3, and −6) was introduced at various positions to probe size constraints and electronegativity, as well as enhance any metabolic instability. Several nitrogen containing bicyclic systems (UP-10 through UP-13) were introduced to probe the size of the binding site. The ketone (UP-7) and benzyl ether (UP-8) were designed to probe the size of the binding site and a potential hydrogen bond with R74, and the aniline (UP-9), which was likely more soluble, tested whether polarity or electronegativity contributed to binding.

**Table 1.**
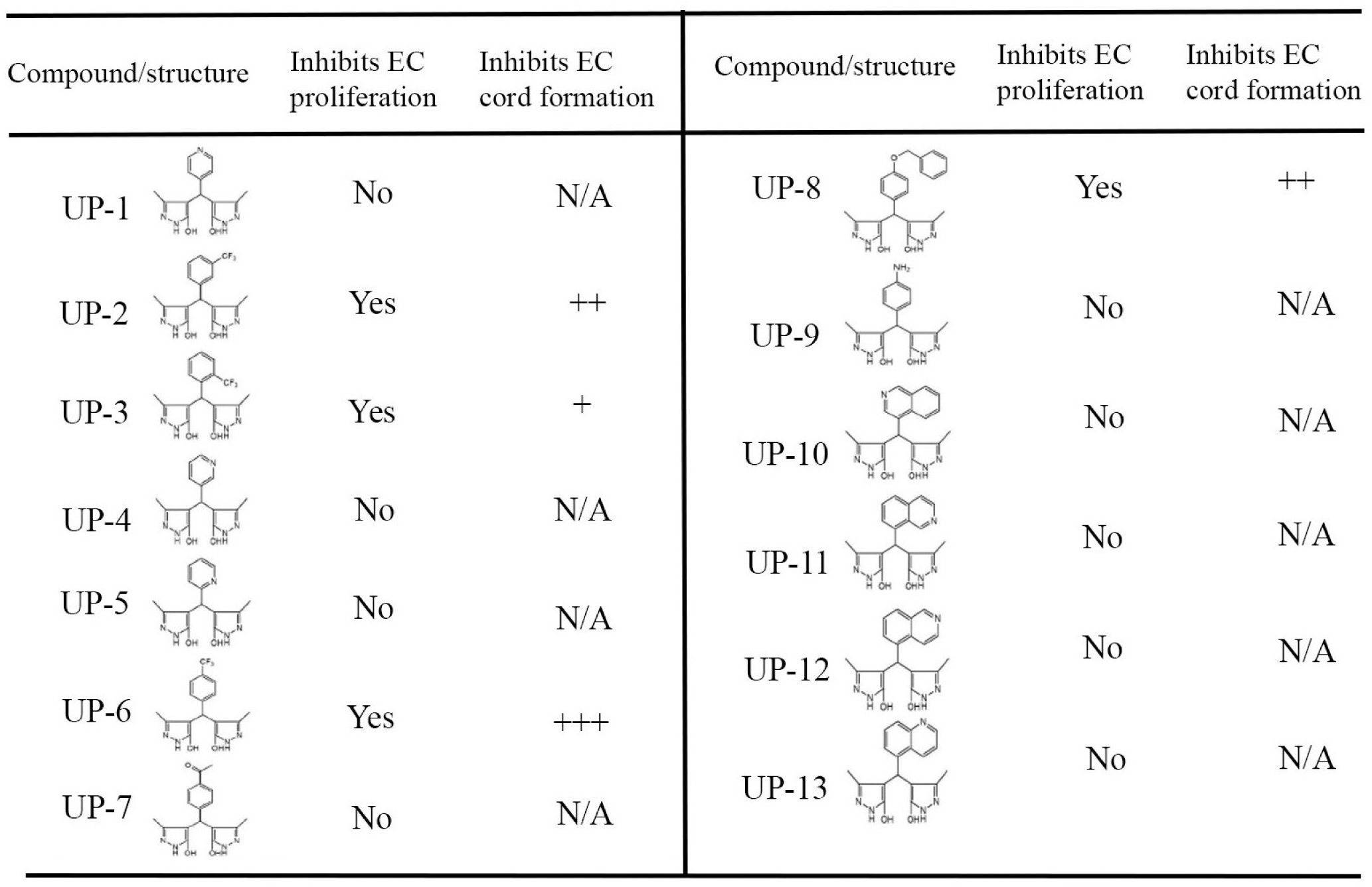
Summary of initial screening results of chemically synthesized C74 analogs. Structures of analogs and their impact on EC proliferation and/or follow-on cord formation assay. Only those compounds which inhibited EC proliferation relative to DMSO treatment control were subjected to cord formation assay for anti-angiogenic activity (+++ indicates highly effective in inhibiting angiogenesis at a 20 μM concentration; N/A – not applicable).

**Gau et al. Table 1**
| Compound/structure | Inhibits EC proliferation | Inhibits EC cord formation | Compound/structure | Inhibits EC proliferation | Inhibits EC cord formation |
| --- | --- | --- | --- | --- | --- |
| UP-1 | No | N/A | UP-8 | Yes | ++ |
| UP-2 | Yes | ++ | UP-9 | No | N/A |
| UP-3 | Yes | + | UP-10 | No | N/A |
| UP-4 | No | N/A | UP-11 | No | N/A |
| UP-5 | No | N/A | UP-12 | No | N/A |
| UP-6 | Yes | +++ | UP-13 | No | N/A |
| UP-7 | No | N/A |  |  |  |

We performed an initial testing of these compounds in EC proliferation assays at a single dose of 20 μM, and then selected those compounds (UP-2, −3, −6 and −8) that inhibited EC proliferation relative to DMSO control for qualitative evaluation of anti-angiogenic activity in EC cord formation assay. Based on this initial qualitative screening, we identified compound UP-6 (4,4’-(4-(trifluoromethyl)phenyl)methylene)bis(3,5-dimethyl-1*H*-pyrazole)) with predicted docked structure shown in **Figure 3B** to be the most effective analog in terms of its ability to inhibit cord morphogenesis (an indicator of angiogenic activity) of ECs *in vitro* (**Table 1).** Encouragingly, 20 μM UP-6 treatment led to near-complete blockade of endothelial cord formation (**Figures 3C-D),** a feature we had previously observed in ECs when subjected to a much higher concentration (50 μM) of C74 treatment (21). In the same study, we also showed that C74 when applied at 10 μM is completely ineffective in reducing cord formation of EC. Encouragingly, at a 10 μM concentration, UP-6 was still effective in inhibiting EC cord formation (by ∼50%) while C74 was not (**Figures 3E-F**). These data demonstrate that UP-6 outperforms C74 in inhibiting angiogenesis *in vitro*, prompting us to further examine whether this is also true in an *in vivo* setting. Therefore, we performed matrigel plug angiogenesis assay where we co-injected bFGF (a stimulator of angiogenesis)-supplemented growth factor-reduced matrigel with either C74 or UP-6 (at 200 μM) or equivalent DMSO control bilaterally into the flanks of C57/Bl6 mice. CD31 immunostaining of plugs harvested 10 days after injection revealed that treatment of either C74 or UP-6 significantly reduced EC infiltration into the plugs, but the reduction in CD31-positivity of plugs treated with UP-6 was more prominent (>50% decrease relative to control) than that achieved by the treatment of C74 compound (∼25% decrease vs control), and this difference was statistically significant (**Figures 3G-H**). These data suggest that UP-6 has a more potent anti-angiogenic activity than C74, providing the first *in vivo* proof-of-concept that anti-angiogenic efficacy based on inhibition of the Pfn1-actin interaction can be optimized through medicinal chemistry strategies.

To gain insight into why UP-6 biologically outperforms C74, we first queried ChemAxon to compare the predicted solubility and lipophilicity values of these two compounds. Not surprisingly, the predicted solubility and lipophilicity values of C74 and UP-6 (LogS = −3.38 (for C74) and = −3.39 (for UP-6); LogP =1.32 (for C74) and =1.43 (for UP6); S = solubility, P = permeability) were found to be nearly identical. Based on the limited number of analogs tested and the results to date, since we cannot yet formulate a basis for this difference in activity among analogs with similar electronic, size, and solubility/lipophilicity characteristics, we first performed an SPR assay to determine whether UP6 interacts with Pfn1 with much stronger affinity relative to C74. Interestingly, in striking contrast to our observations with C74, UP6-Pfn1 SPR assay revealed inverted (negative) sensorgrams in a dose-dependent fashion (**Figure 4A**). Although not very common, these types of unconventional negative sensorgrams have been previously reported in the literature for certain protein-ligand interactions (33–37). Although the exact structural basis for negative sensorgrams is debatable and not fully understood, major conformational change in the protein upon ligand binding has been hypothesized to be one such reason (35). In fact, when we performed a Differential Scanning Fluorometry (DSF) assay comparing the thermal stability of the Pfn1 protein with compound UP-6 and C74 vs DMSO control, interestingly, both C74 and UP-6 produced negative thermal shifts of Pfn1 relative to DMSO control, but the extent of the negative thermal shift was more pronounced for UP-6 than C74 (**Figure 4B)**. Ligand binding can either stabilize or destabilize protein structure, and the latter is reflected as diminished protein melting temperature upon ligand binding (38). Therefore, our results could mean that these small molecule (C74 or UP-6) interactions might be promoting a less stable Pfn1 conformation, altering dynamics and/or irreversibility of Pfn1 unfolding, or inducing a conformational change that facilitates SYPRO fluorescence at lower temperatures with UP-6 having a more pronounced impact than C74.

**Figure 4.**
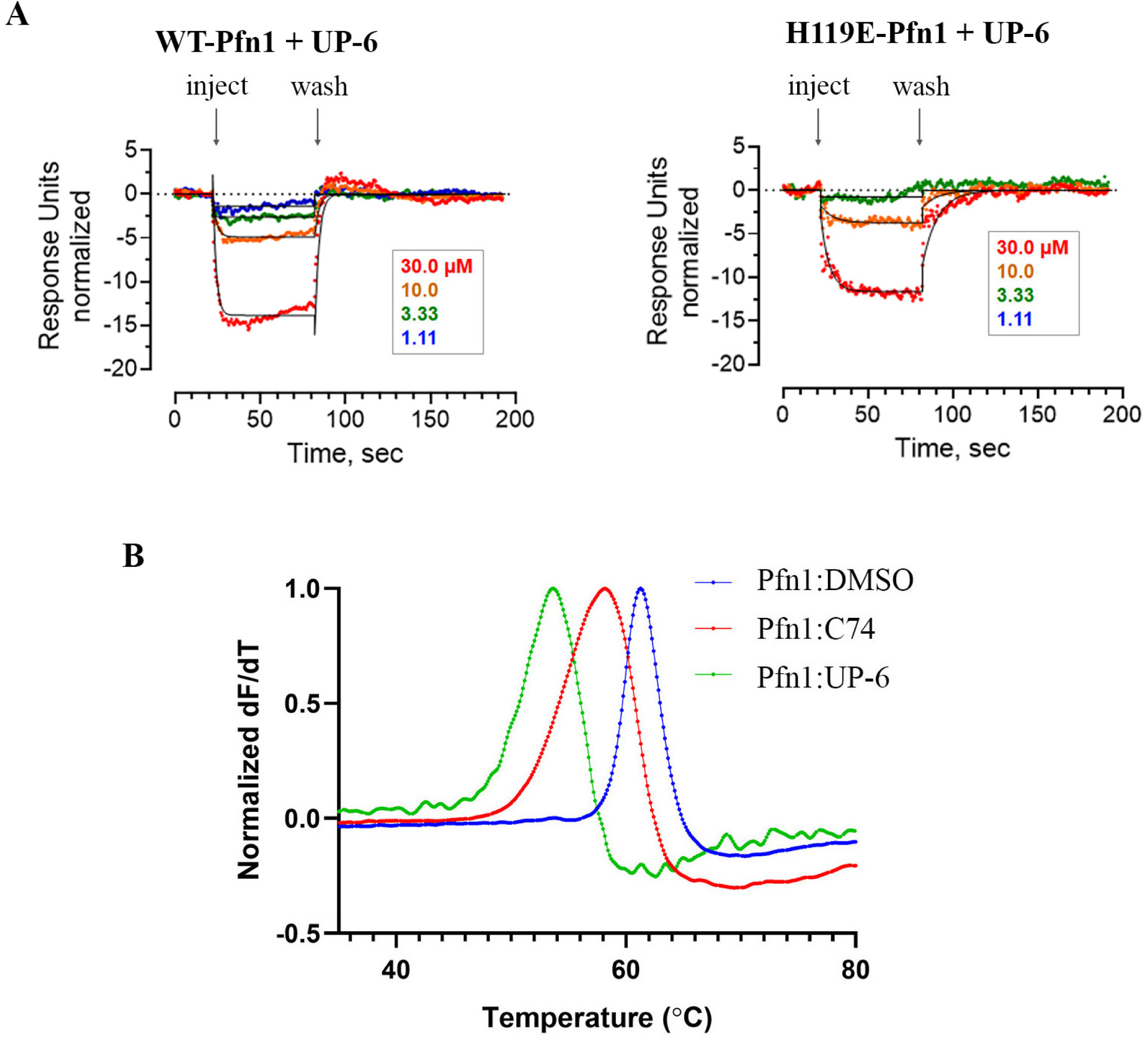
Biochemical characterization of UP-6:Pfn1 interaction by SPR and DSF assays. **A)** Inverted sensorgrams of UP-6-Pfn1 (WT or H119E) interaction as revealed by the SPR assay (the experimental details of the SPR assays are identical to those described for C74 in the legends of Figure 2). **B)** Normalized rate of change in SYPRO fluorescence as a function of temperature from the DSF assay comparing Pfn1 treated with 1mM C74, UP-6, or a DMSO control. Each data point is the average of three technical replicates and normalized to the maximum value within the experiment.

### Feasibility of ultrasound-guided tumor-localized delivery of Pfn1 inhibitor to achieve anti-angiogenic and therapeutic benefit

Unlike subcutaneous tumors, access to internal organ tumors (such as RCC) is challenging for direct injection, limiting the feasibility of localized administration. Furthermore, although low-affinity inhibitors can minimize prolonged off-target toxicity, global systemic delivery of low-affinity inhibitors may require high doses to achieve therapeutic concentrations at the tumor site, potentially leading to off-target effects and toxicity. Since Pfn1 is ubiquitously expressed and a central regulator of actin cytoskeleton irrespective of cell type, there could be additional challenges associated with global systemic delivery of high doses of Pfn1 inhibitor for therapeutic applications. To overcome the key limitations associated with systemic and direct injection approaches, we wanted to explore whether Pfn1 inhibitor can be systemically released in a tumor-localized manner to achieve therapeutic benefit. Lipid microbubbles serve as efficient carriers of various agents (small molecules and nucleic acids) and can release their payloads for localized therapeutic applications when subjected to ultrasound-targeted microbubble destruction (UTMD)(39–42). The other advantages of this platform are: a) the utility of microbubbles as ultrasound contrast agents for *in vivo* imaging, and b) the rapid clearance of microbubbles from circulation through hepatic and splenic filtration, with ∼95% cleared from bloodstream within 30 minutes (43). First, as a demonstration of feasibility, we were able to successfully encapsulate C74 within lipid microbubbles (∼3 µm in diameter), as confirmed by MS evaluation of microbubbles with estimated concentration of 0.190 ng/µL (see **Supplementary Information** *Section E*). Second, as a proof-of-concept *in vitro* study, we performed HmVEC cord morphogenesis assay where we demonstrated that similar to the action of unencapsulated C74 (a positive control), UTMD of C74-encapsulated microbubbles prominently inhibits cord morphogenesis of ECs (**Figures 5**). The extent of angiogenesis inhibition was more pronounced when C74 was released from microbubbles compared to the setting where C74 was directly added to the culture. We speculate that this could be due to the possible concentration differences of C74 between these two settings. To establish the specificity of our findings, we included several negative control groups in that same experimental setting where either C74-encapsulated microbubbles was not subjected to UTMD or C74 was transiently released by UTMD followed by immediate C74 washout and replacement with regular culture media or UTMD of microbubbles was performed without C74 as a payload (to rule out any possible detrimental effect of ultrasound cavitation on cells). In none of these negative control settings did we observe any adverse effect on endothelial cord formation when compared to the control DMSO treatment group.

**Figure 5.**
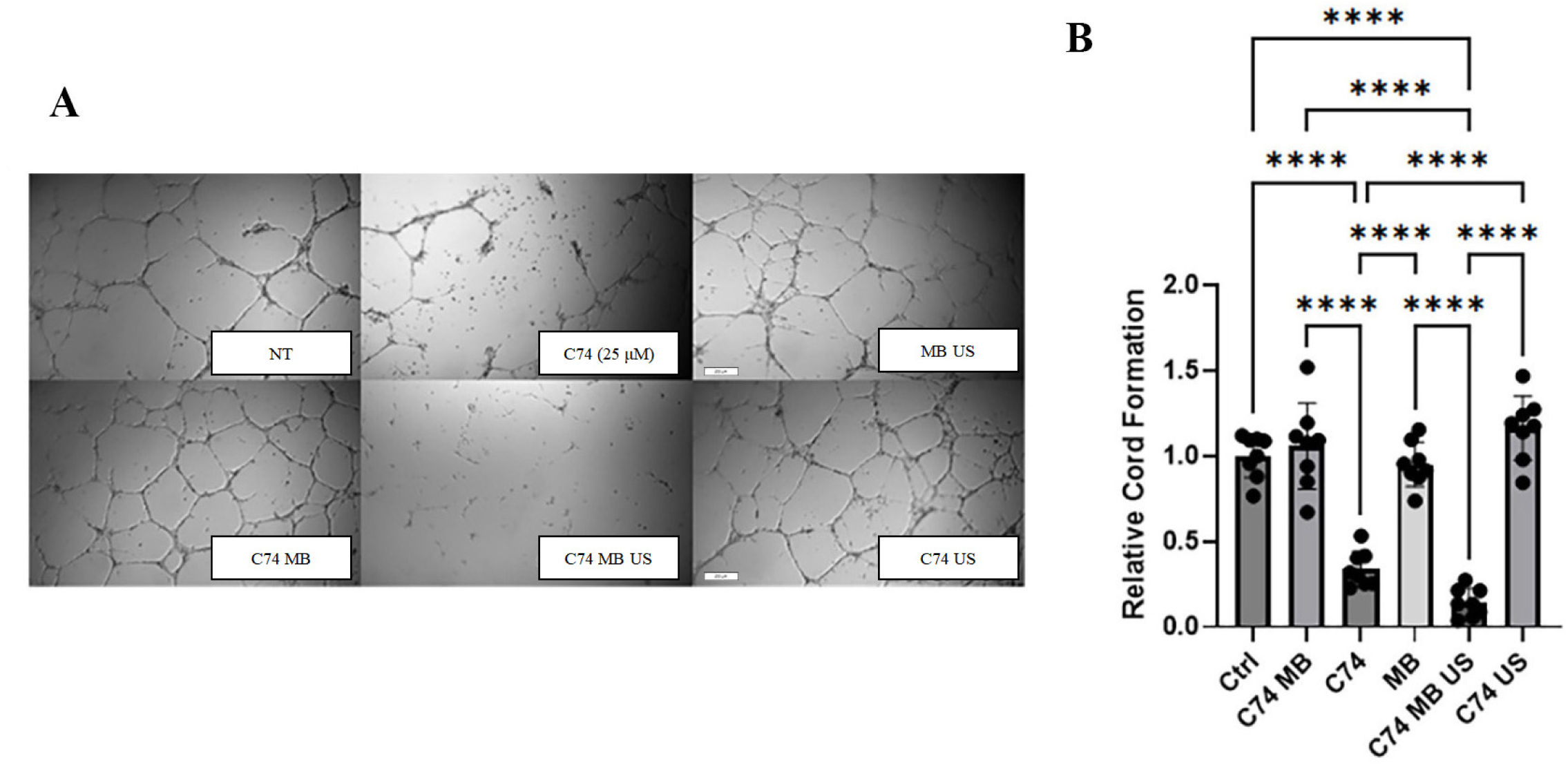
*In vitro* proof-of-concept of angiogenesis inhibition by C74 upon release by UTMD of lipid microbubbles. **A-B)** Representative cord-formation images of HmVECs for various treatment settings (*panel A*): NT: DMSO (vehicle control); C74: treatment with unencapsulated C74 (this group shows effect of C74 alone); MB US: empty MB with UTMD (this control group tests if bubble cavitation alone causes reduction in cord formation), C74 MB: C74-encapsulated MB with no UTMD (this group demonstrates that un-cavitated bubbles do not cause an effect even though they are dosed with C74); C74 MB US: C74-encapsulated MB with UTMD (this is the true test group showing that when bubbles are destroyed and C74 is released, we see reduction in cord formation), and C74 US: C74-encapsulated MB with UTMD followed by immediate washout and replacement with C74-devoid culture media (this group tests whether C74 immediately enters cells post-bubble destruction). The bar graph in *panel B* depicts the relative cord lengths between the different treatment groups - cord length is defined as the distance between the junctional i.e. cord-intersectional points. (One-way ANOVA with Tukey post-hoc test, **** p <0.0001 from n = 3 experiments).

Given these results demonstrating the feasibility of UTMD to release C74 and achieve anti-angiogenic action in cell culture settings, we next performed exploratory *in vivo* tumor model studies. Specifically, we subcutaneously implanted RENCA cells in Balb/c mice to establish tumors. After allowing these tumors to reach palpable size, tumor-bearing mice received retro-orbital injections of C74-loaded microbubbles three times over the course of a week (with a 100 μl injection of microbubble solution per treatment, this is equivalent to administration of ∼190 ng of C74 administration per injection), followed by ultrasound application to target the tumor site (the experimental schema is shown in **Figure 6A**; **Figure 6B** shows the ultrasound images of microbubbles at the tumor site before and after UTMD). Consistent with our in vitro findings, this precision delivery approach led to a significant ∼30% reduction in the average tumor burden and a concomitant decrease in tumor angiogenesis (**Figures 6C-F**). We also extended these *in vivo* studies with UP-6-loaded microbubbles (as per MS evaluation, the estimated concentration of UP6 in microbubble solution (=0.221 ng/ µL; **Supplemental Information** *Section E*) was nearly comparable to that estimated for C74-loaded microbubbles (=0.19 ng/µL)). With a 100 μl injection of microbubble solution per treatment, this is equivalent to administration of 221 ng of UP6 per treatment. Although we have no way of accurately estimating the exact amount of each compound released in the tumor microenvironment due to possible experiment-to-experiment as well as animal-to-animal variations in cavitation efficiency and clearing of microbubbles, encouragingly UTMD of UP6-loaded microbubbles also diminished tumor angiogenesis and led to a greater inhibition (∼50% reduction) in the average tumor burden than we observed in the C74 setting (**Figures 6G-J**). Collectively, these results support the utility of lipid microbubble-based delivery of Pfn1 inhibitor as a promising targeted therapy to diminish neovascularization and tumor growth.

**Figure 6:**
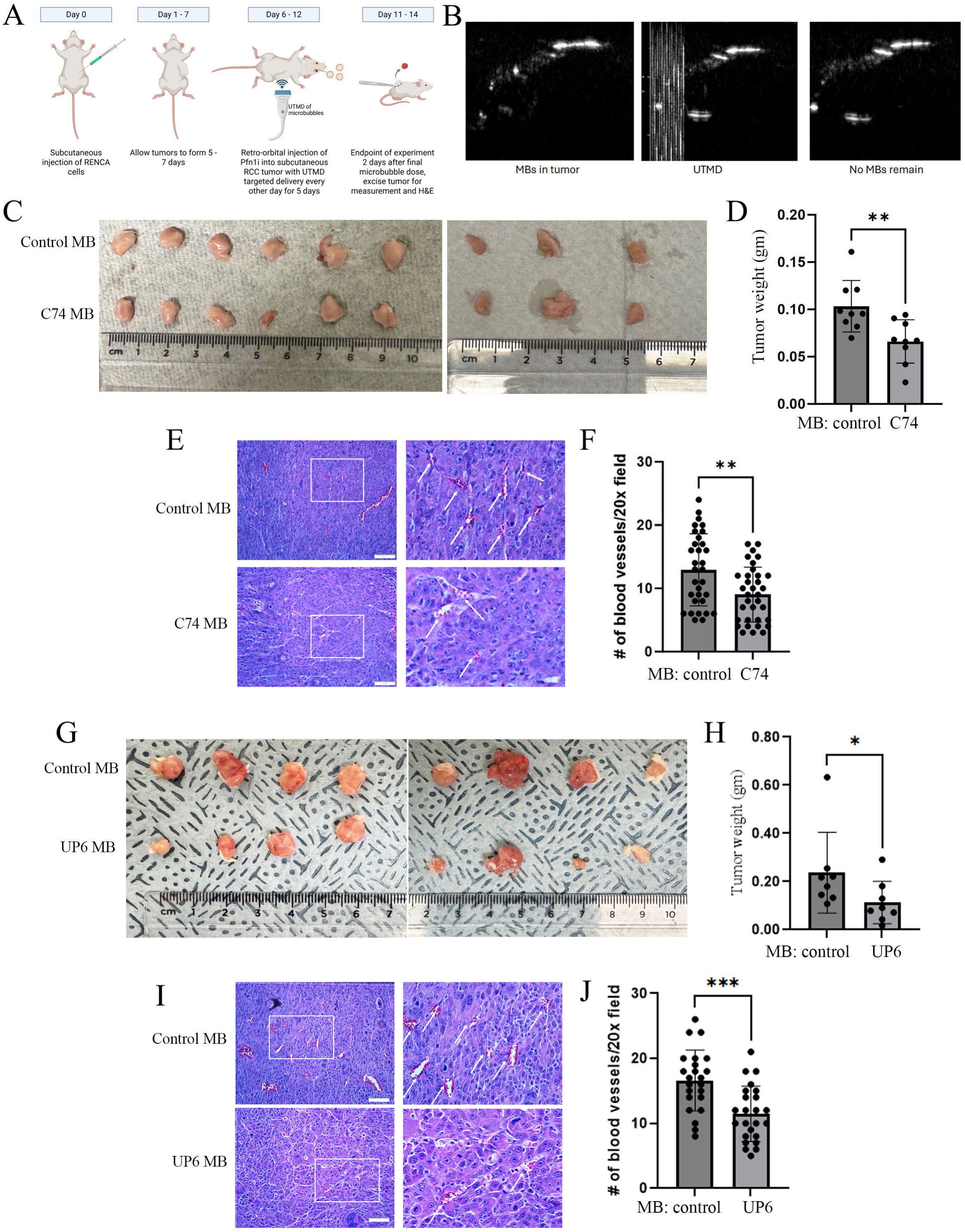
Feasibility of ultrasound-guided tumor-localized delivery of Pfn1 inhibitor to achieve anti-angiogenic and therapeutic benefit. **A)** Experimental schema of tumor microbubble studies (MB – microbubbles). **B)** Ultrasound images (*panel B*) showing MB infiltration into tumor vasculature (white arrows), followed by UTMD and no contrast remaining in the tumor space. **C-J)** Results of C74-MB (*panels C through F*) and UP6-MB (*panels G-J*) tumor model studies. Images of subcutaneous RENCA tumors subjected to UTMD of indicated MBs (*panels C, G*) with end-point tumor burden comparison between the control and experimental treatment groups (*panels D and H*). Representative H&E images of tumor histosections (*panels E and I;* white arrows indicate functional tumor-associated blood vessels with presence of red blood cells) and the associated quantification (*panels F and I*) of the average number of tumor-associated blood vessels per 20X field of observation between the two treatment groups (Mann-Whitney test, *: p<0.05; ** p < 0.01, ***: p<0.001; data pooled from 8-9 mice per group; scale bar, 50 μm).

## DISCUSSION

In our previous studies, we established genetic proof-of-concept for a critical requirement for endothelial Pfn1 activity for developmental (9) as well as tumor angiogenesis (15). While our previously identified small molecule inhibitors of the Pfn1-actin interaction exhibited anti-angiogenic activity in ocular settings(9, 21), given that tumor angiogenesis involves complex interplay of multiple different cell types in the TME, the overall angiogenic impact of small molecules targeting Pfn1’s action in a complex, multicellular TME remained an open question. In this investigation, we report four new findings to substantially advance our previous studies. First, our SPR studies show that our previously identified small molecule inhibitor of Pfn1-actin interaction, C74, directly binds to Pfn1. Second, this study provides the first demonstration of small molecule intervention of Pfn1 as a strategy to reduce tumor angiogenesis in RCC. Third, we were able to design and chemically synthesize a new structural analog of C74 (UP-6) with improved anti-angiogenic activity *in vitro* as well as *in vivo*. Fourth, we provide the first proof-of-concept for the feasibility of ultrasound-mediated disruption of circulating lipid microbubbles for tumor-localized delivery of Pfn1 inhibitors achieve anti-angiogenic and anti-tumor benefit *in vivo*.

In the RENCA tumor model, the degree to which tumor angiogenesis was inhibited by C74 was less than we had previously seen in the setting of genetic disruption of Pfn1 in ECs (15). This can be due to several reasons. First, our SPR studies predicted that C74 is a low-affinity inhibitor of Pfn1 with a dissociation constant measured to be approximately 60 μM (a value that is in general agreement with our previously published data where C74 treatment in a 25 – 50 μM concentration range effectively inhibited RCC cell migration/proliferation and angiogenic activity of ECs). Second, small molecules released in the TME are expected to be taken up by multiple different types of cells, and therefore, its effect is not restricted to ECs only. For example, fibroblasts and various immune cells also contribute to tumor angiogenesis (44, 45), and at present, we do not have a clear understanding of how functional inhibition of Pfn1 impacts angiogenesis-promoting activities of those cell types in the TME. Third, since Pfn1 also interacts with many other proteins besides actin, small molecule-mediated inhibition of Pfn1-actin interaction cannot be equated to complete genetic loss-of-function of Pfn1. Fourth, we cannot rule out off-target effects of C74 that can potentially offset the anti-angiogenic effect of the inhibition on Pfn1-actin interaction. Endothelial transcriptomic studies comparing the impact of genetic loss-of-function vs small molecule inhibition may provide greater insights into the on-target specificity of C74’s action.

Although our proof-of-concept tumor model studies were conducted in a subcutaneous tumor setting (a limitation of this study), our ability to attenuate angiogenesis and tumor reduction upon UTMD of systemically circulating microbubbles sets the stage for future exploration in an orthotopic RCC tumor setting to better model the disease. Our findings support the therapeutic potential of lipid microbubble-based delivery systems as a clinically translatable strategy for targeted cancer therapy. Notably, microbubbles are already FDA-approved as contrast agents for ultrasound imaging (Definity, SonoVue), which provides a strong foundation for their repurposing in therapeutic applications. In recent years, microbubble-assisted delivery has gained traction in oncology, with studies demonstrating enhanced delivery of chemotherapeutics, nucleic acids, and small molecules to solid tumors, including pancreatic (46–48), liver (49, 50), and brain (51) tumors, when combined with UTMD. This platform enables site-specific release of therapeutic payloads, reducing systemic toxicity while enhancing local efficacy. Our study adds to this growing body of evidence by showing that microbubble-mediated delivery of Pfn1 inhibitor can diminish tumor growth and angiogenesis in RCC. These data highlight the promise of leveraging microbubble technology for precision delivery of anti-tumor agents, particularly for tumors located in anatomically challenging or sensitive regions. Furthermore, given our initial indication for improved anti-angiogenic activity of compound UP-6 vs C74, we will need to further explore the anti-tumor efficacy of compound UP6 in preclinical orthotopic settings of RCC, either as a stand-alone agent or in combination with other exiting anti-angiogenic drugs.

Another limitation of the present study is that the exact underlying basis for improved biological activity of compound UP-6 over C74 is not clear. Although we do not have direct experimental evidence, given that the predicted solubility and lipophilicity values of these two compounds are nearly equal, we speculate that differential cell permeability is unlikely to be the reason for an improved biological activity of UP6. The structure activity relationship (**Table 1**) suggests that modifications of the aryl group affect binding. This may be due to changes in the electronic structure of the ring changing the favorability of interactions with R74 and H119 (**Figure 3**). However, C74 binding is maintained in the H119E mutant (**Figure 2D**), which is unexpected if this histidine is making a cation-pi or halogen bond. This may indicate the histidine interaction is not important or that there are inaccuracies in the docking model. The negative thermal shift (**Figure 4B**) supports that there may be a significant conformational change in the protein that is not accounted for in the docking model, which used a rigid protein structure. Dose-dependent inverted sensorgram characteristics of UP6:Pfn1 interaction in our SPR assay (**Figure 4A**) as well as a greater thermal shift of Pfn1 by compound UP-6 relative to C74 in the DSF assay (**Figure 4B**) could be indicative of an even larger structural alteration in Pfn1 elicited by its interaction with UP-6. Although it is beyond the scope of the present work, X-ray crystallography studies in the future are expected to provide valuable structural insights into how these small molecule interactions impact the atomic arrangements in the Pfn1 molecule.

Finally, the therapeutic utility of targeting the Pfn1-actin interaction extends beyond cancer. Excessive or abnormal angiogenesis is also a hallmark of several other diseases, including age-related macular degeneration, diabetic retinopathy, atherosclerosis, and psoriasis. In some of these conditions, unchecked angiogenesis leads to the formation of dysfunctional blood vessels that contribute to disease progression and tissue damage. Given the clinical evidence for endothelial Pfn1 upregulation in at least atherosclerosis and diabetic retinopathy(9, 52–56), small molecule inhibition of Pfn1 could be a worthwhile strategy to explore for those disease conditions in the future.

## ACKNOWLEDGMENT

The authors acknowledge funding from NIH grants R01CA248873 (Roy), R01CA271095 (Roy), R21EY-032632 (Roy, Huryn, Koes, and VanDemark), K99/R00 CA267189 (Gau), T32-HL129964 (Villanueva) and Department of Defense grants HT425-24-2-0556 (Roy) and W81XWH-19-1-0768 (Roy).

## AUTHOR CONTRIBUTIONS

DG performed experiments, conceived study design, analyzed data, wrote the manuscript, and acquired funding; PC, KX, NW, JA, CV, MW, SZ and MW performed experiments and/or generated reagents; CC generated reagents and wrote the manuscript; HS performed SPR experiments and wrote the manuscript; XC, AP, and FV assisted with and oversaw microbubble studies; TES oversaw the SPR studies and wrote the manuscript; PF performed molecular docking studies; DK, AV, DMH, and PR conceived study design, wrote the manuscript, and acquired funding.

## DATA AVAILABILITY

All data are included in either main figures or supplementary information (SI).

## Notes

**CONFLICT OF INTEREST:** The authors declare no conflict of interest.

### Competing Interest Statement

The authors have declared no competing interest.

### Summary of Updates

Addition of new data and reorganization of text

