## Supplementary Info for "Small molecule intervention of actin-binding protein profilin1 reduces tumor angiogenesis in renal cell carcinoma"

**Supplementary information guide:**

Section A - General Information,

Section B - General procedure of compound synthesis

Section C - Compound information

Section D - NMR data of compounds

Section E - Mass-spectrometry data of lipid microbubbles loaded with compounds (C74 and UP6)

### Section A - General Information

C74 analogs were either purchased from commercial sources or chemically synthesized. Reagents and solvents used in chemical synthesis were purchased from commercial sources at the highest commercial quality. Thin layer chromatography (TLC) was performed using Millipore Sigma™ silica gel 60 F254 coated aluminum-backed TLC sheets. TLC spots were detected using 254 nm UV light. $^1$H NMR was used to confirm the purity of all the final products. Deuterated dimethylsulfoxide (DMSO-$d_6$) was used as NMR solvent. $^1$H NMR and $^{13}$C NMR spectra were recorded on Bruker AVANCE NEO 400 or 600 MHz spectrometers. Chemical shifts (δ) are reported in parts per million (ppm) relative to internal residual solvent peaks from indicated deuterated solvents. Coupling constants (J) are reported in Hertz (Hz) and are rounded to the nearest 0.1 Hz. Multiplicities are defined as: s = singlet, d = doublet, t = triplet, q = quartet, m = multiplet, dd = doublet of doublets, dt = doublet of triplets, ddd = doublet of doublet of doublets, dddd = doublet of doublet of doublet of doublets, br = broad, app = apparent, par = partial. Accurate mass measurement experiments were performed using direct infusion electrospray ionization (ESI) or atmospheric pressure photoionization (APPI) on a Bruker scimaX 7T Fourier-transform ion cyclotron resonance mass spectrometer. Molecular formulae are confirmed using the SmartFormulaReport plugin in the Compass Data Analysis software. The tolerated error for molecular formula confirmation is no more than 3 ppm.

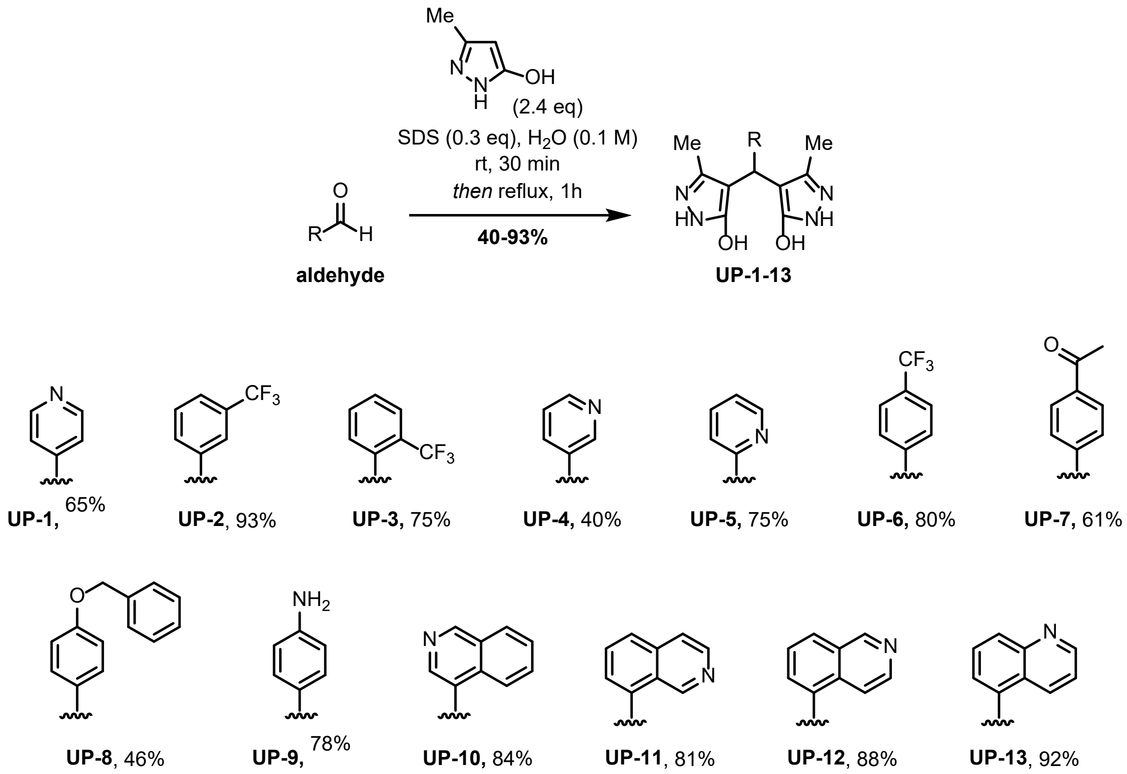


**Figure 1.** Scheme for the synthesis of UP-1 through UP-13 (**3a-m**).

### Section B - General procedure for synthesis

To a round-bottom flask was added corresponding **aldehyde** (1.0 eq), **3-methyl-5-pyrazolone** (2.4 eq), sodium dodecyl sulfate (0.3 eq), and $H_2O$ (0.1 M). The reaction mixture was stirred at room temperature for 30 minutes and then refluxed until starting **aldehyde** appeared consumed by TLC (10% methanol in DCM). The precipitate that was formed over the course of the reaction was filtered off and washed with $H_2O$ (3 x 15 mL), then dried by stirring while heating to 80 °C under vacuum overnight to give the product as a solid.

### Section C - Compound Information (UP compounds 1-13)

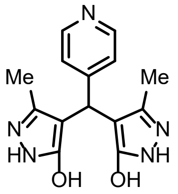

**UP-1**

**UP-1**: 4,4'-(pyridin-4-ylmethylene)bis(3-methyl-1H-pyrazol-5-ol). **pink solid, 65%.**

AMM (ESI) m/z: $[M + H]^+$ Calcd for $C_{14}H_{16}N_5O_2$ 286.129851; found 286.130129.

$^1$H NMR (400 MHz, DMSO-$d_6$) δ 11.31 (s, 4H), 8.43 – 8.37 (m, 2H), 7.11 (d, *J* = 5.1 Hz, 2H), 4.85 (s, 1H), 2.10 (s, 6H).

$^{13}$C{$^1$H} NMR (151 MHz, DMSO-$d_6$) δ 160.98, 152.36, 149.06, 139.92, 123.09, 103.02, 32.35, 10.35.

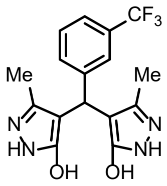

**UP-2**

**UP-2:** 4,4'-((3-(trifluoromethyl)phenyl)methylene)bis(3-methyl-1H-pyrazol-5-ol). **peach solid, 93%.**

AMM (ESI) m/z: $[M + H]^+$ Calcd for $C_{16}H_{16}F_3N_4O_2$ 353.121987; found 353.121947.

$^1$H NMR (600 MHz, DMSO-$d_6$) δ 11.38 (s, 4H), 7.52 – 7.40 (m, 4H), 4.94 (s, 1H), 2.09 (s, 6H).

$^{13}$C{$^1$H} NMR (151 MHz, DMSO-$d_6$) δ 161.01, 144.85, 139.93, 131.89, 128.87, 128.55 (q, *J* = 31.2 Hz), 124.52 (q, *J* = 272.4 Hz), 123.78 (q, *J* = 3.9 Hz), 122.42 (q, *J* = 4.2 Hz), 103.69, 32.66, 10.36.

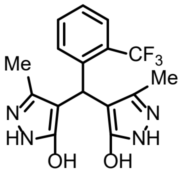

**UP-3**

**UP-3**: 4,4'-((2-(trifluoromethyl)phenyl)methylene)bis(3-methyl-1H-pyrazol-5-ol). **light yellow solid, 75%.**

AMM (ESI) m/z: $[M + H]^+$ Calcd for $C_{16}H_{16}F_3N_4O_2$ 353.121987; found 353.122045.

$^1$H NMR (400 MHz, DMSO-$d_6$) δ 10.71 (s, 4H), 7.74 (d, *J* = 8.0 Hz, 1H), 7.62 (d, *J* = 7.9 Hz, 1H), 7.55 (t, *J* = 7.6 Hz, 1H), 7.36 (t, *J* = 7.5 Hz, 1H), 5.29 (s, 1H), 1.82 (s, 6H).

$^{13}$C{$^1$H} NMR (151 MHz, DMSO-$d_6$) δ 160.14, 142.28, 137.79, 132.07, 131.63, 126.42, 126.36 (q, *J* = 29.3 Hz), 125.71 (q, *J* = 6.0 Hz), 124.81 (q, *J* = 274.6 Hz), 102.95, 30.67, 10.36.

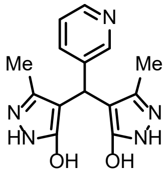

**UP-4**

**UP-4**: 4,4'-(pyridin-3-ylmethylene)bis(3-methyl-1H-pyrazol-5-ol). **pink solid, 40%**.

AMM (ESI) m/z: $[M + H]^+$ Calcd for $C_{14}H_{16}N_5O_2$ 286.129851; found 286.130258.

$^1$H NMR (400 MHz, DMSO-$d_6$) δ 11.39 (s, 4H), 8.33 (dd, *J* = 4.4, 1.8 Hz, 2H), 7.52 (dt, *J* = 8.1, 2.2 Hz, 1H), 7.24 (dd, *J* = 7.8, 4.8 Hz, 1H), 4.89 (s, 1H), 2.10 (s, 6H).

$^{13}$C{$^1$H} NMR (151 MHz, DMSO-$d_6$) δ 160.97, 149.07, 146.61, 139.69, 138.68, 135.02, 122.89, 103.44, 30.70, 10.34.

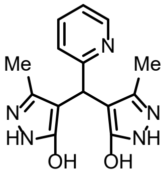

**UP-5**

**UP-5**: 4,4'-(pyridin-2-ylmethylene)bis(3-methyl-1H-pyrazol-5-ol). **peach solid, 75%.**

AMM (ESI) m/z: $[M + H]^+$ Calcd for $C_{14}H_{16}N_5O_2$ 286.129851; found 286.129893.

$^1$H NMR (400 MHz, DMSO-$d_6$) δ 11.25 (s, 4H), 8.43 (dd, *J* = 5.1, 1.8 Hz, 1H), 7.71 (td, *J* = 7.7, 1.8 Hz, 1H), 7.38 (d, *J* = 7.9 Hz, 1H), 7.24 – 7.16 (m, 1H), 4.98 (s, 1H), 2.02 (s, 6H).

$^{13}$C{$^1$H} NMR (151 MHz, DMSO-$d_6$) δ 162.60, 160.31, 147.86, 138.94, 136.91, 122.52, 121.36, 103.15, 36.67, 10.43.

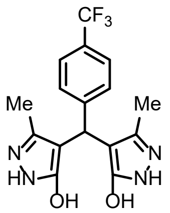

**UP-6**

**UP-6**: 4,4'-((4-(trifluoromethyl)phenyl)methylene)bis(3-methyl-1H-pyrazol-5-ol). **peach solid, 80%**.

AMM (ESI) m/z: $[M + H]^+$ Calcd for $C_{16}H_{16}F_3N_4O_2$ 353.121987; found 353.122049.

$^1$H NMR (600 MHz, DMSO-$d_6$) δ 11.37 (s, 4H), 7.58 (d, *J* = 8.1 Hz, 2H), 7.32 (d, *J* = 8.0 Hz, 2H), 4.91 (s, 1H), 2.09 (s, 6H).

$^{13}$C{$^1$H} NMR (151 MHz, DMSO-$d_6$) δ 160.96, 148.28, 139.82, 128.26, 126.25 (q, *J* = 31.6 Hz), 124.63 (q, *J* = 4.2 Hz), 124.55 (q, *J* = 271.5 Hz), 103.62, 32.72, 10.32.

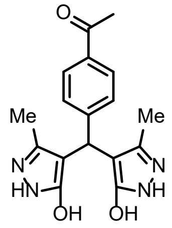

**UP-7**

**UP-7**: 1-(4-(bis(5-hydroxy-3-methyl-1H-pyrazol-4-yl)methyl)phenyl)ethan-1-one. **pink solid, 61%.**

AMM (ESI) m/z: $[M + H]^+$ Calcd for $C_{17}H_{19}N_4O_3$ 327.145167; found 327.145362.

$^1$H NMR (400 MHz, DMSO-$d_6$) δ 11.33 (s, 4H), 7.85 – 7.78 (m, 2H), 7.25 (d, *J* = 8.2 Hz, 2H), 4.89 (s, 1H), 2.52 (s, 3H), 2.08 (s, 6H).

$^{13}$C{$^1$H} NMR (151 MHz, DMSO-$d_6$) δ 197.62, 161.03, 149.18, 139.82, 134.53, 127.93, 127.82, 103.75, 32.97, 26.69, 10.41.

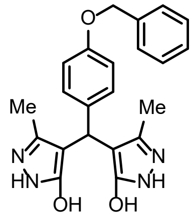

**UP-8**

**UP-8**: 4,4'-((4-(benzyloxy)phenyl)methylene)bis(3-methyl-1H-pyrazol-5-ol). **yellow solid, 46%.**

AMM (ESI) m/z: $[M + H]^+$ Calcd for $C_{22}H_{23}N_4O_3$ 391.176467; found 391.176442.

$^1$H NMR (600 MHz, DMSO-$d_6$) δ 11.33 (s, 4H), 7.42 (d, *J* = 7.5 Hz, 2H), 7.37 (t, *J* = 7.5 Hz, 2H), 7.31 (t, *J* = 7.3 Hz, 1H), 7.03 (d, *J* = 8.3 Hz, 2H), 6.85 (d, *J* = 8.4 Hz, 2H), 5.04 (s, 2H), 4.76 (s, 1H), 2.07 (s, 6H).

$^{13}$C{$^1$H} NMR (151 MHz, DMSO-$d_6$) δ 161.59, 156.76, 140.14, 137.80, 135.99, 128.87, 128.20, 128.07, 114.49, 104.98, 69.61, 32.44, 10.84.

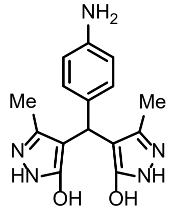

**UP-9**

**UP-9**: 4,4'-((4-aminophenyl)methylene)bis(3-methyl-1H-pyrazol-5-ol). **yellow solid, 78%.**

AMM (ESI) m/z: $[M + H]^+$ Calcd for $C_{15}H_{18}N_5O_2$ 300.145501; found 300.145699.

$^1$H NMR (400 MHz, DMSO-$d_6$) δ 11.23 (s, 4H), 6.80 – 6.74 (m, 2H), 6.43 – 6.37 (m, 2H), 4.76 (s, 1H), 4.64 (s, 1H), 2.04 (s, 6H).

$^{13}$C{$^1$H} NMR (151 MHz, DMSO-$d_6$) δ 161.17, 146.15, 139.60, 130.48, 127.87, 113.49, 104.95, 31.91, 10.43.

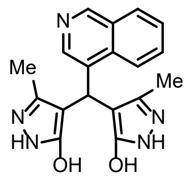

**UP-10**

**UP-10**: 4,4'-(isoquinolin-4-ylmethylene)bis(3-methyl-1H-pyrazol-5-ol). **peach solid, 84%.**

AMM (ESI) m/z: $[M + H]^+$ Calcd for $C_{18}H_{18}N_5O_2$ 336.145501; found 336.145603.

$^1$H NMR (600 MHz, DMSO-$d_6$) δ 10.56 (d, *J* = 160.5 Hz, 4H), 9.15 (s, 1H), 8.26 (s, 1H), 8.10 (dd, *J* = 8.4, 1.3 Hz, 1H), 7.86 (d, *J* = 8.5 Hz, 1H), 7.71 (ddd, *J* = 8.4, 6.8, 1.4 Hz, 1H), 7.62 (ddd, *J* = 8.0, 6.7, 1.1 Hz, 1H), 5.55 (s, 1H), 1.80 (s, 6H).

$^{13}$C{$^1$H} NMR (151 MHz, DMSO-$d_6$) δ 159.95, 151.01, 142.40, 137.64, 133.64, 131.36, 130.14, 128.15, 127.94, 126.65, 122.90, 102.26, 28.76, 10.42.

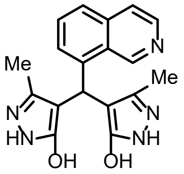



UP-11

**UP-11**: 4,4'-(isoquinolin-8-ylmethylene)bis(3-methyl-1H-pyrazol-5-ol). **peach solid, 81%.**

AMM (ESI) m/z: $[M + H]^+$ Calcd for $C_{18}H_{18}N_5O_2$ 336.145501; found 336.145608.

$^1H$ NMR (400 MHz, DMSO-$d_6$) δ 10.60 (s, 4H), 9.28 (s, 1H), 8.44 (d, *J* = 5.6 Hz, 1H), 7.82 – 7.76 (m, 2H), 7.65 (t, *J* = 7.7 Hz, 1H), 7.39 (d, *J* = 7.1 Hz, 1H), 5.78 (s, 1H), 1.74 (s, 6H).

$^{13}C\{^1H\}$ NMR (151 MHz, DMSO-$d_6$) δ 159.82, 148.90, 142.11, 140.02, 137.65, 135.92, 129.76, 126.93, 126.04, 125.10, 120.89, 102.76, 30.09, 10.41.

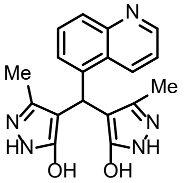



UP-12

**UP-12**: 4,4'-(isoquinolin-5-ylmethylene)bis(3-methyl-1H-pyrazol-5-ol). **peach solid, 88%.**

AMM (ESI) m/z: $[M + H]^+$ Calcd for $C_{18}H_{18}N_5O_2$ 336.145501; found 336.145553.

$^1H$ NMR (400 MHz, DMSO-$d_6$) δ 10.59 (s, 4H), 9.27 (s, 1H), 8.42 (d, *J* = 6.0 Hz, 1H), 7.94 (dd, *J* = 7.5, 1.8 Hz, 1H), 7.67 (d, *J* = 6.0 Hz, 1H), 7.62 – 7.51 (m, 2H), 5.56 (s, 1H), 1.75 (s, 6H).

$^{13}C\{^1H\}$ NMR (151 MHz, DMSO-$d_6$) δ 159.93, 152.91, 142.62, 138.29, 137.65, 133.67, 129.62, 128.71, 126.63, 125.97, 116.83, 102.48, 30.27, 10.42.

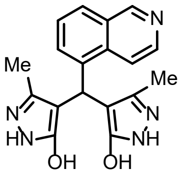



UP-13

**UP-13**: 4,4'-(quinolin-5-ylmethylene)bis(3-methyl-1H-pyrazol-5-ol). **white solid, 92%.**

AMM (ESI) m/z: $[M + H]^+$ Calcd for $C_{18}H_{18}N_5O_2$ 336.145501; found 336.145530.

$^1H$ NMR (400 MHz, DMSO-$d_6$) δ 10.58 (s, 4H), 8.84 (d, *J* = 4.0 Hz, 1H), 8.26 (d, *J* = 8.6 Hz, 1H), 7.86 (d, *J* = 8.3 Hz, 1H), 7.63 (t, *J* = 7.8 Hz, 1H), 7.45 (dd, *J* = 8.6, 4.2 Hz, 1H), 7.36 (d, *J* = 7.2 Hz, 1H), 5.61 (s, 1H), 1.71 (s, 6H).

$^{13}C\{^1H\}$ NMR (151 MHz, DMSO-$d_6$) δ 159.85, 149.66, 148.32, 139.74, 137.64, 132.13, 128.58, 127.51, 126.18, 125.91, 120.82, 102.68, 30.33, 10.42.

### Section - NMR Data

$^{1}H$ NMR spectrum of **UP-1** (400 MHz, DMSO-$d_6$)

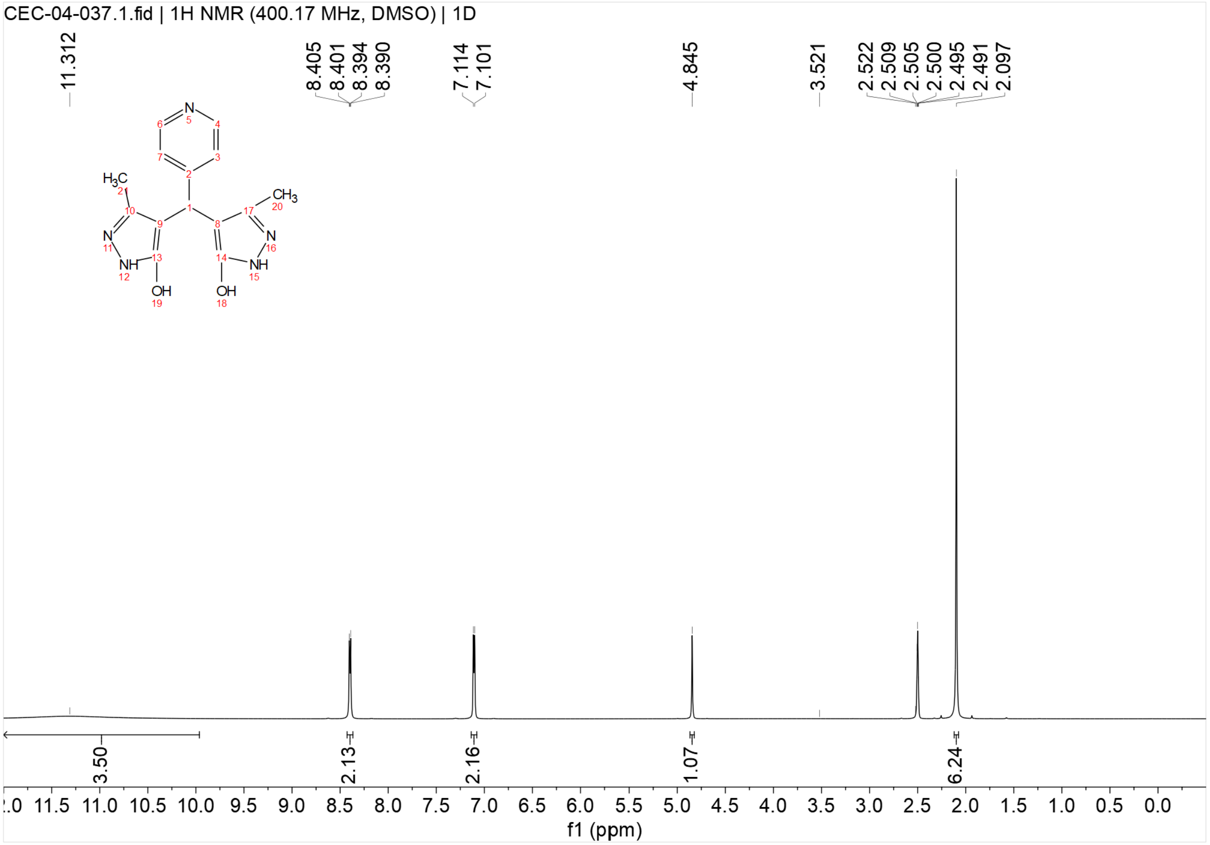


$^{13}C\{^{1}H\}$ NMR spectrum of **UP-1** (151 MHz, DMSO-$d_6$)

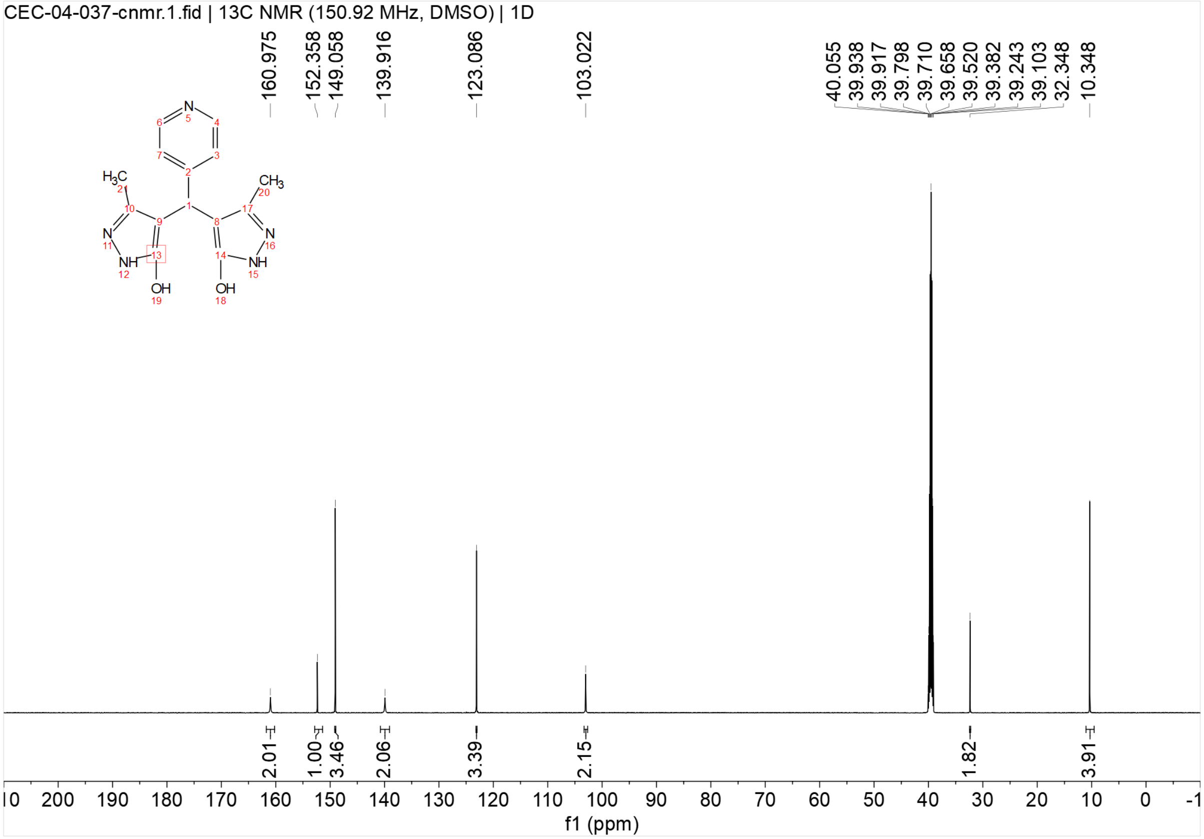
CEC-04-037-cnmr.1.fid | 13C NMR (150.92 MHz, DMSO) | 1D
160.975
152.358
149.058
139.916
123.086
103.022
40.055
39.938
39.917
39.798
39.710
39.658
39.520
39.382
39.243
39.103
32.348
10.348
2.01
1.00
3.46
2.06
3.39
2.15
1.82
3.91
f1 (ppm)

$^{1}H$ NMR spectrum of **UP-2** (600 MHz, DMSO-$d_6$)

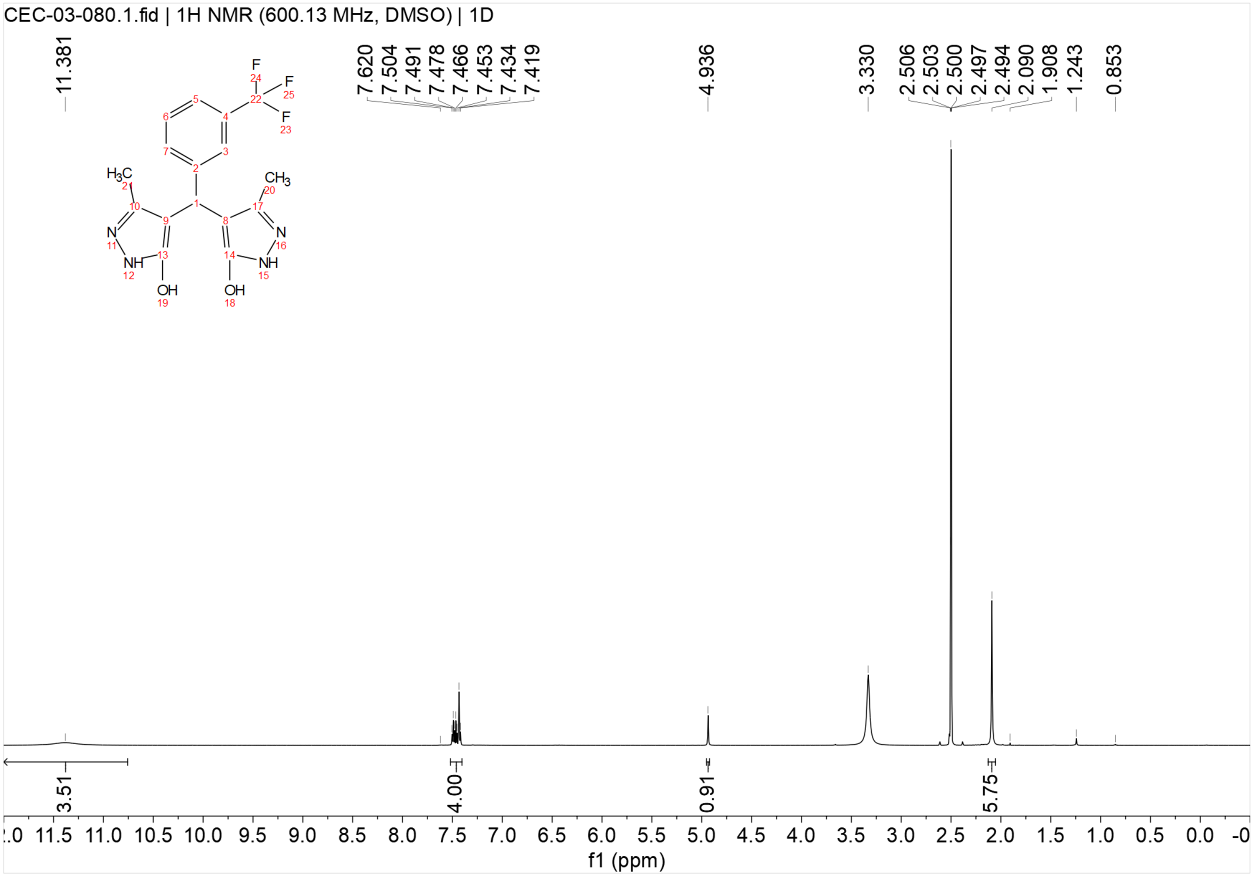


$^{13}C\{^{1}H\}$ NMR spectrum of **UP-2** (151 MHz, DMSO-$d_6$)

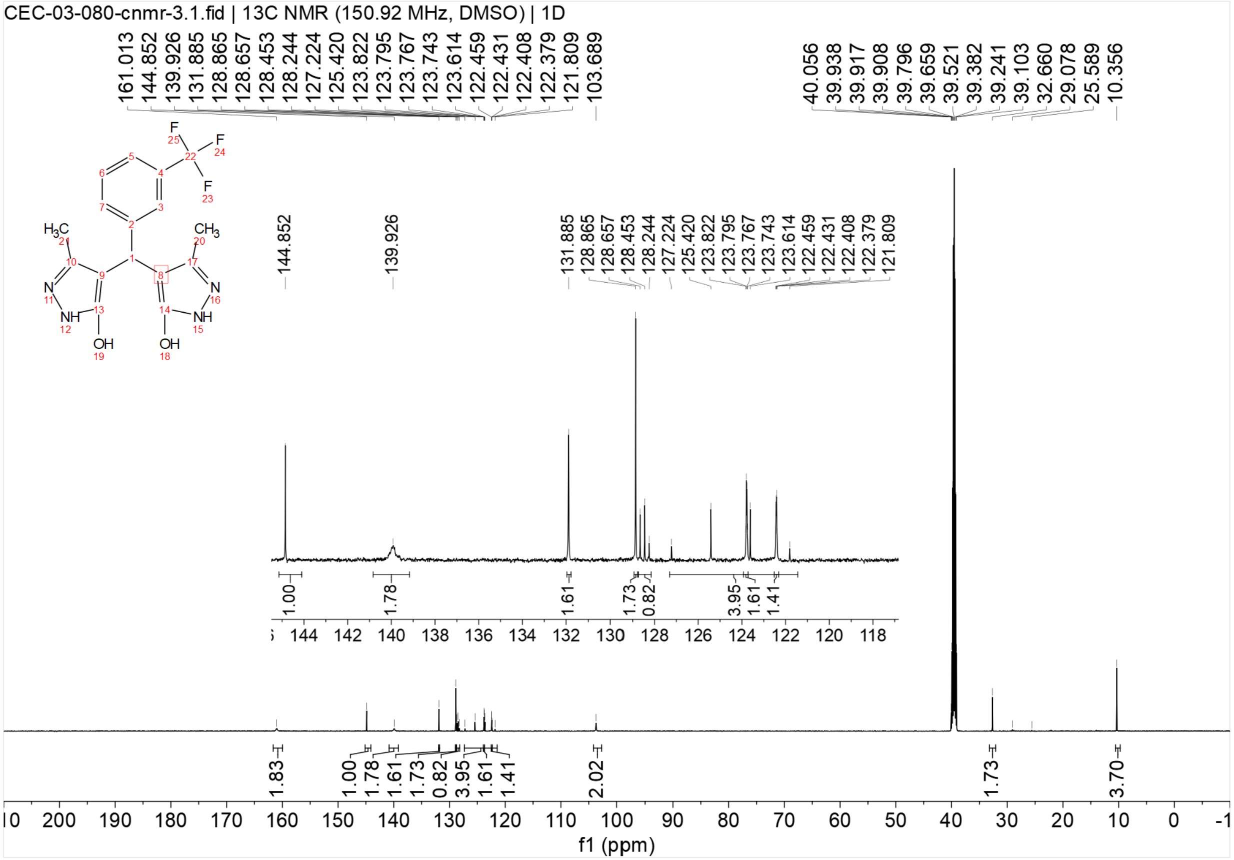


[1]H NMR spectrum of **UP-3** (400 MHz, DMSO-$d_6$)

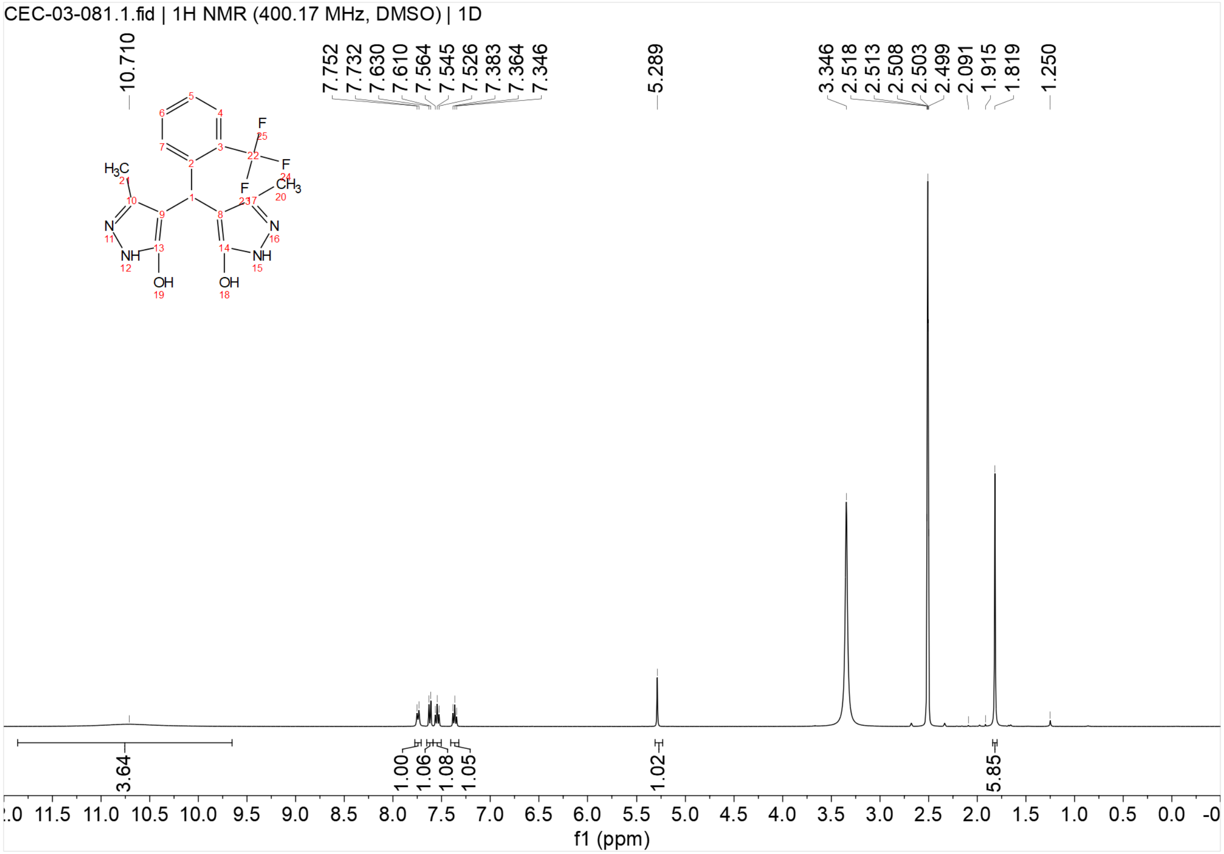

$^{13}C\{^{1}H\}$ NMR spectrum of **UP-3** (151 MHz, DMSO-$d_6$)

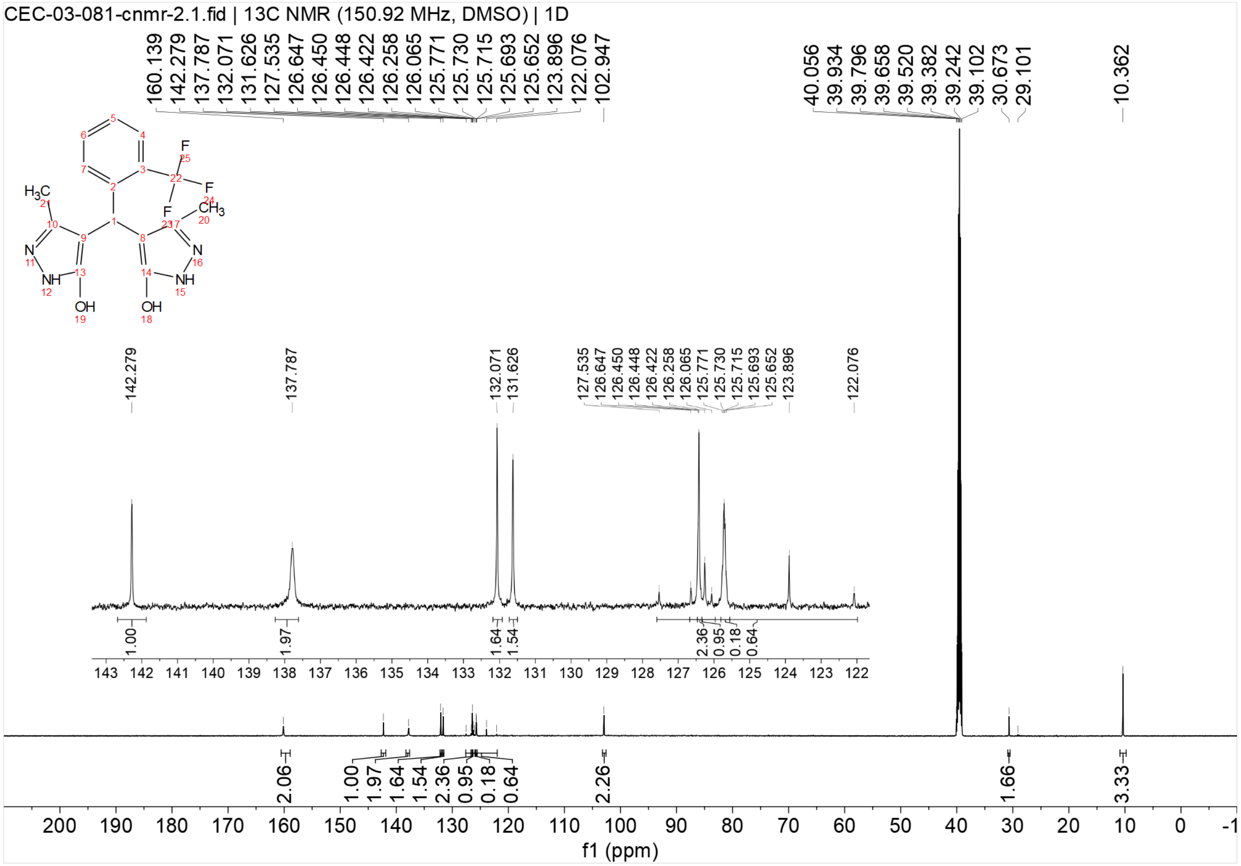


$^{1}H$ NMR spectrum of **UP-4** (400 MHz, DMSO-$d_6$)

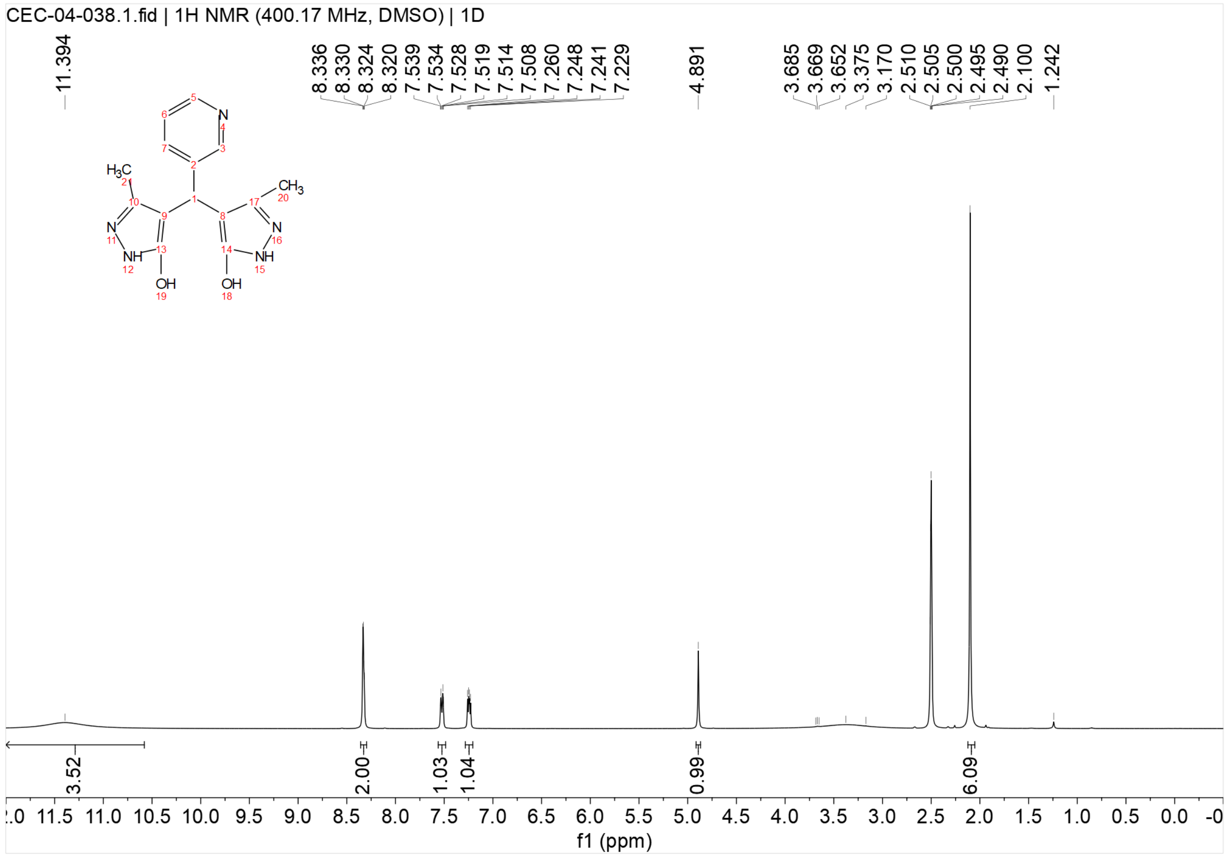


$^{13}C\{^{1}H\}$ NMR spectrum of **UP-4** (151 MHz, DMSO-$d_6$)

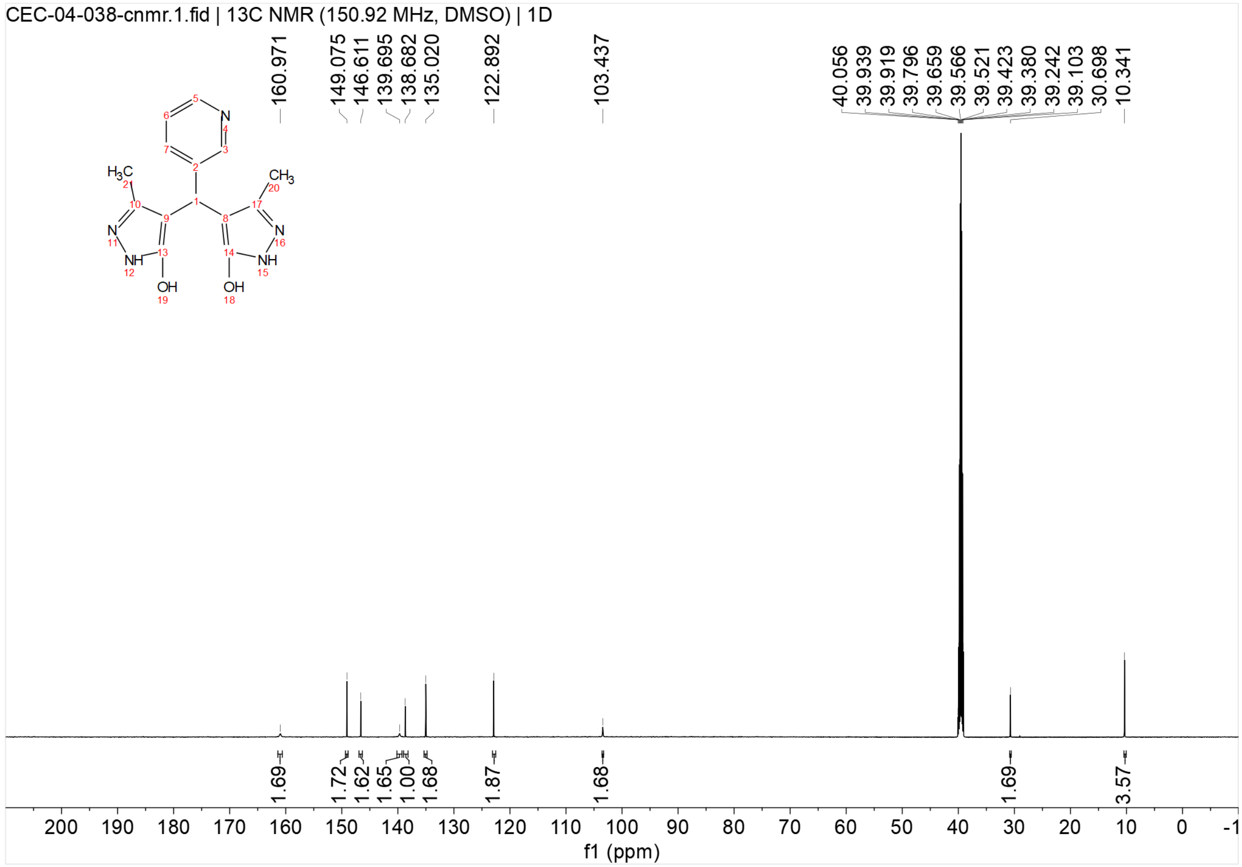

[1]H NMR spectrum of **UP-5** (400 MHz, DMSO-$d_6$)

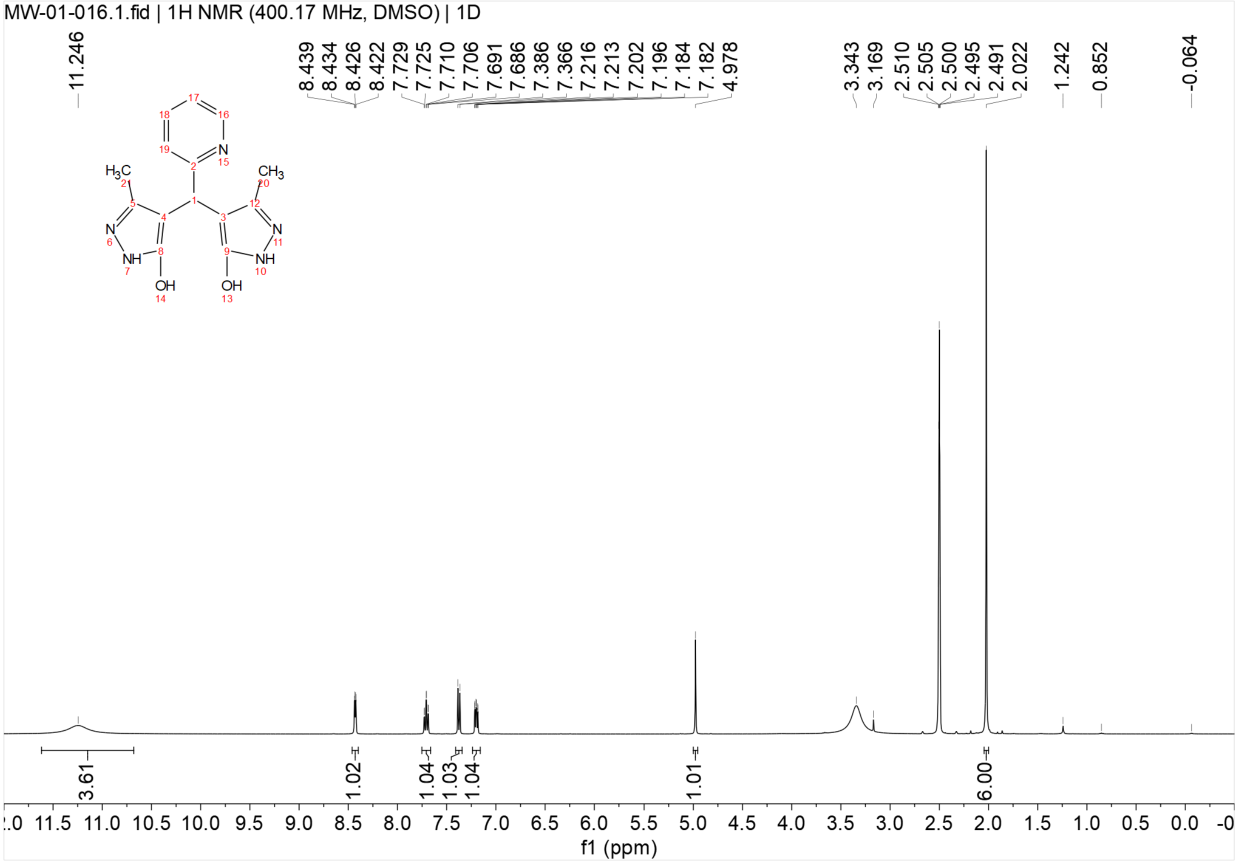


[13]C{[1]H} NMR spectrum of **UP-5** (151 MHz, DMSO-$d_6$)

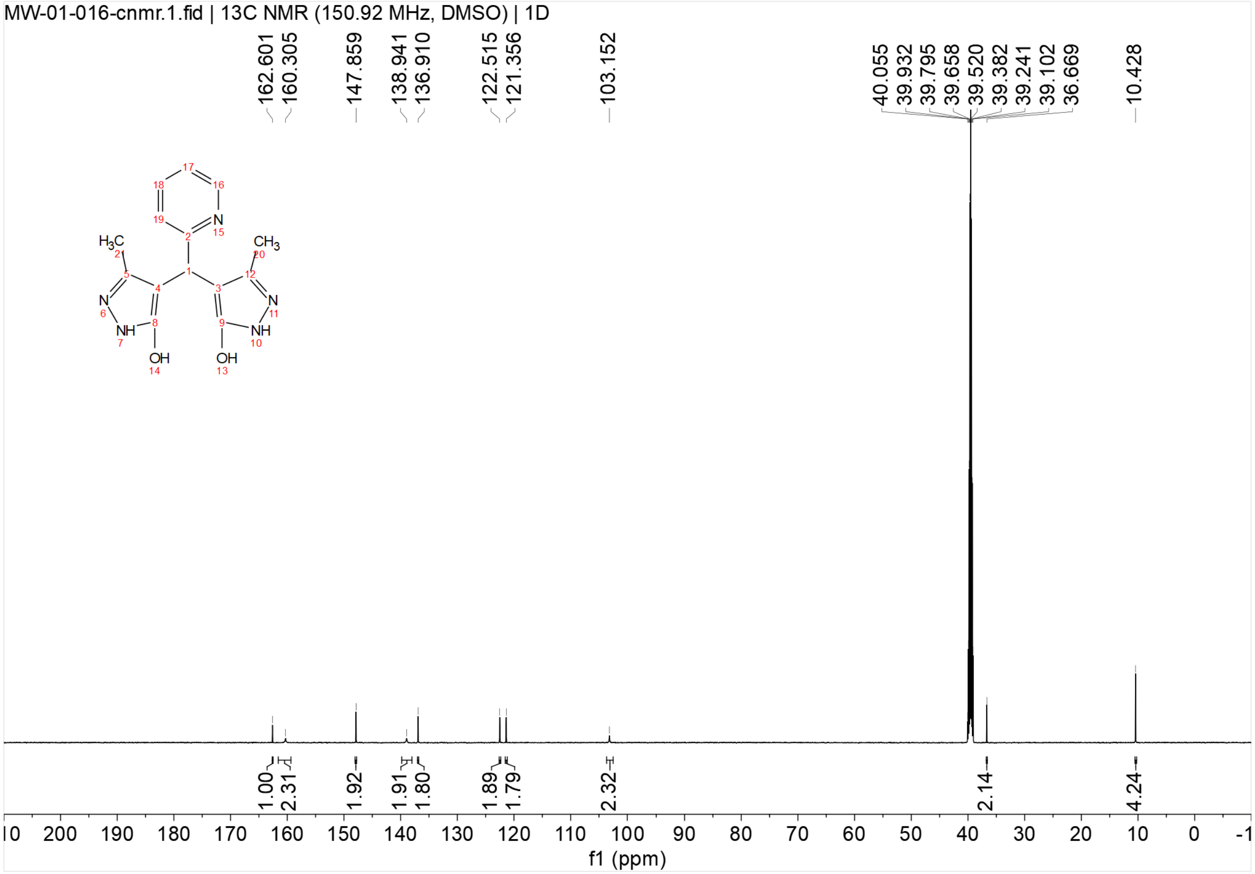


$^1$H NMR spectrum of **UP-6** (600 MHz, DMSO-$d_6$)

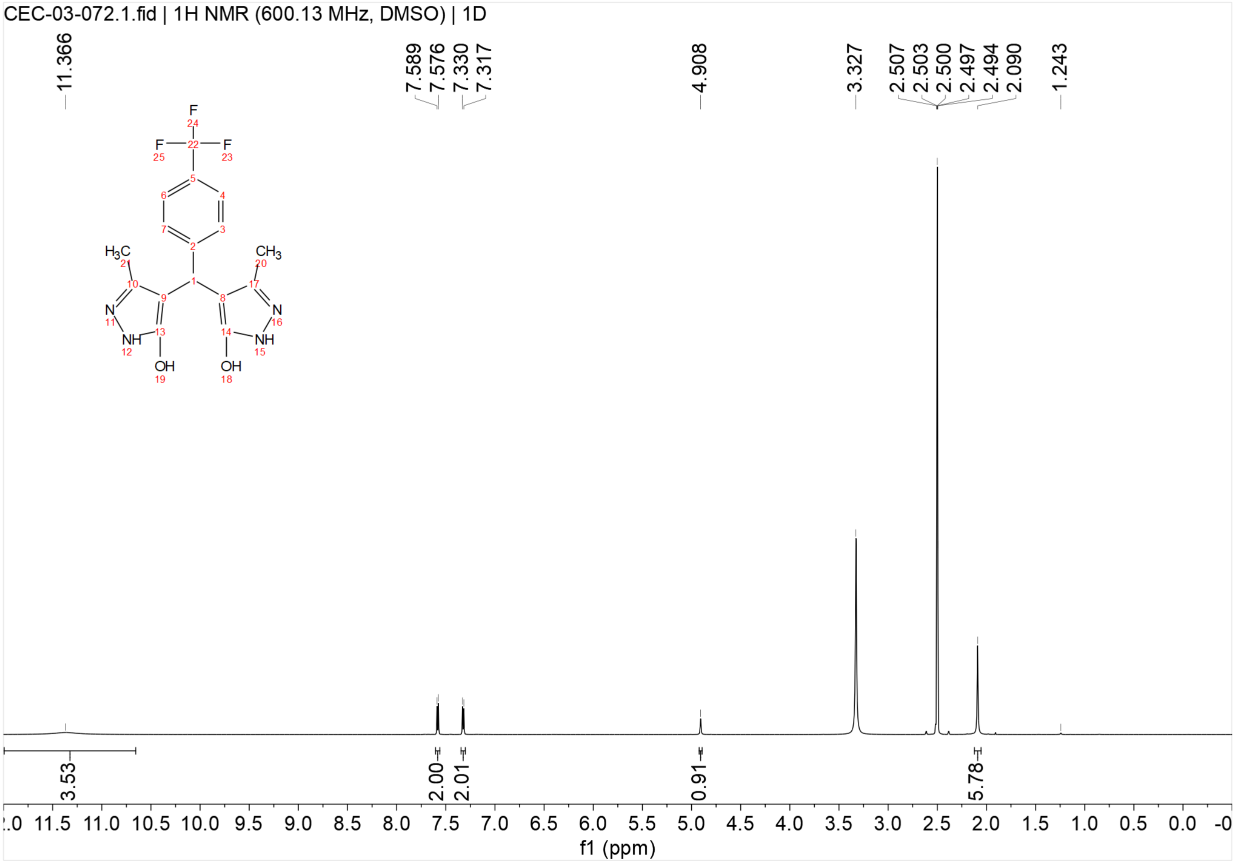


$^{13}C\{^{1}H\}$ NMR spectrum of **UP-6** (151 MHz, DMSO-$d_6$)

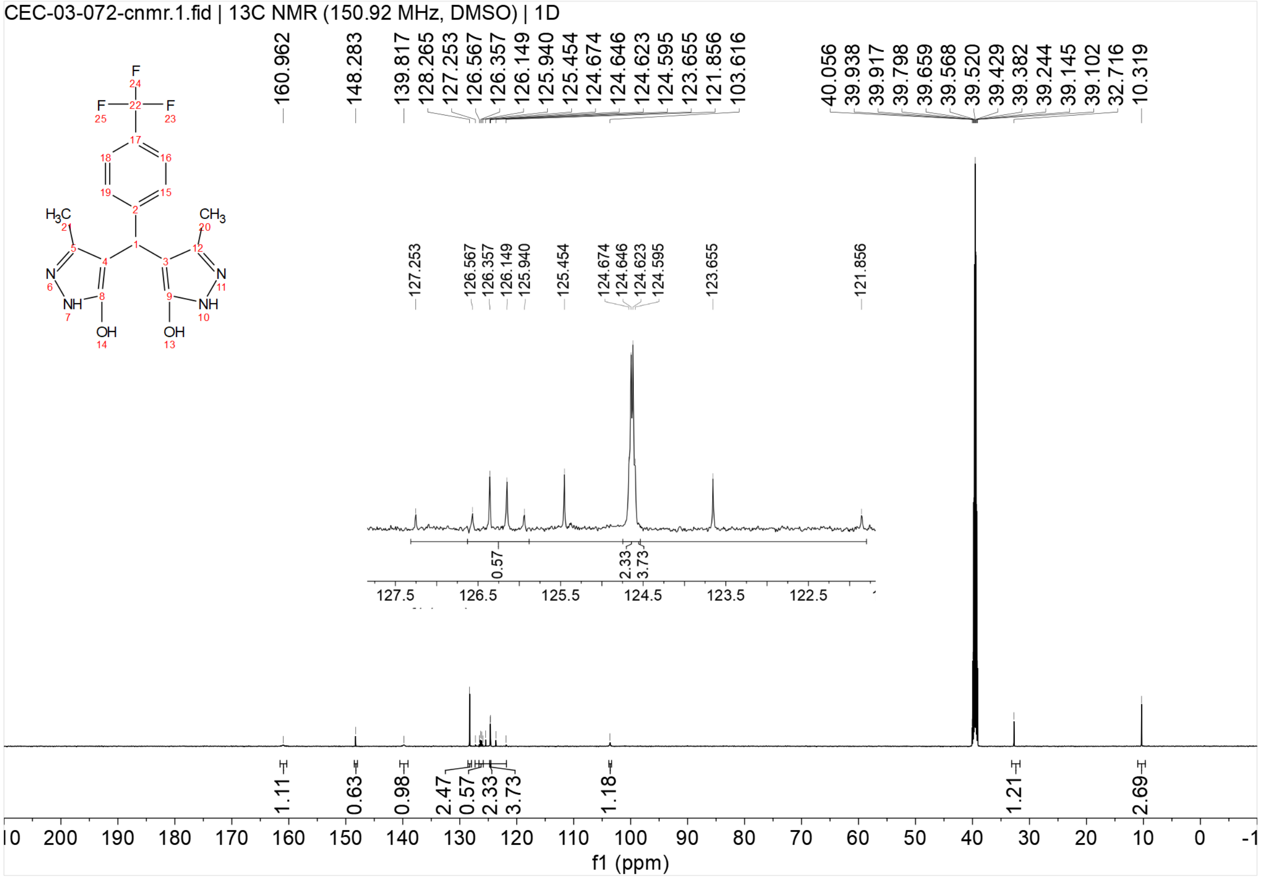
CEC-03-072-cnmr.1.fid | 13C NMR (150.92 MHz, DMSO) | 1D
160.962
148.283
139.817
128.265
127.253
126.567
126.357
126.149
125.940
125.454
124.674
124.646
124.623
124.595
123.655
121.856
103.616
40.056
39.938
39.917
39.798
39.659
39.568
39.520
39.429
39.382
39.244
39.145
39.102
32.716
10.319
1.11
0.63
0.98
2.47
0.57
2.33
3.73
1.18
1.21
2.69
f1 (ppm)

[1]H NMR spectrum of **UP-7** (400 MHz, DMSO-$d_6$)

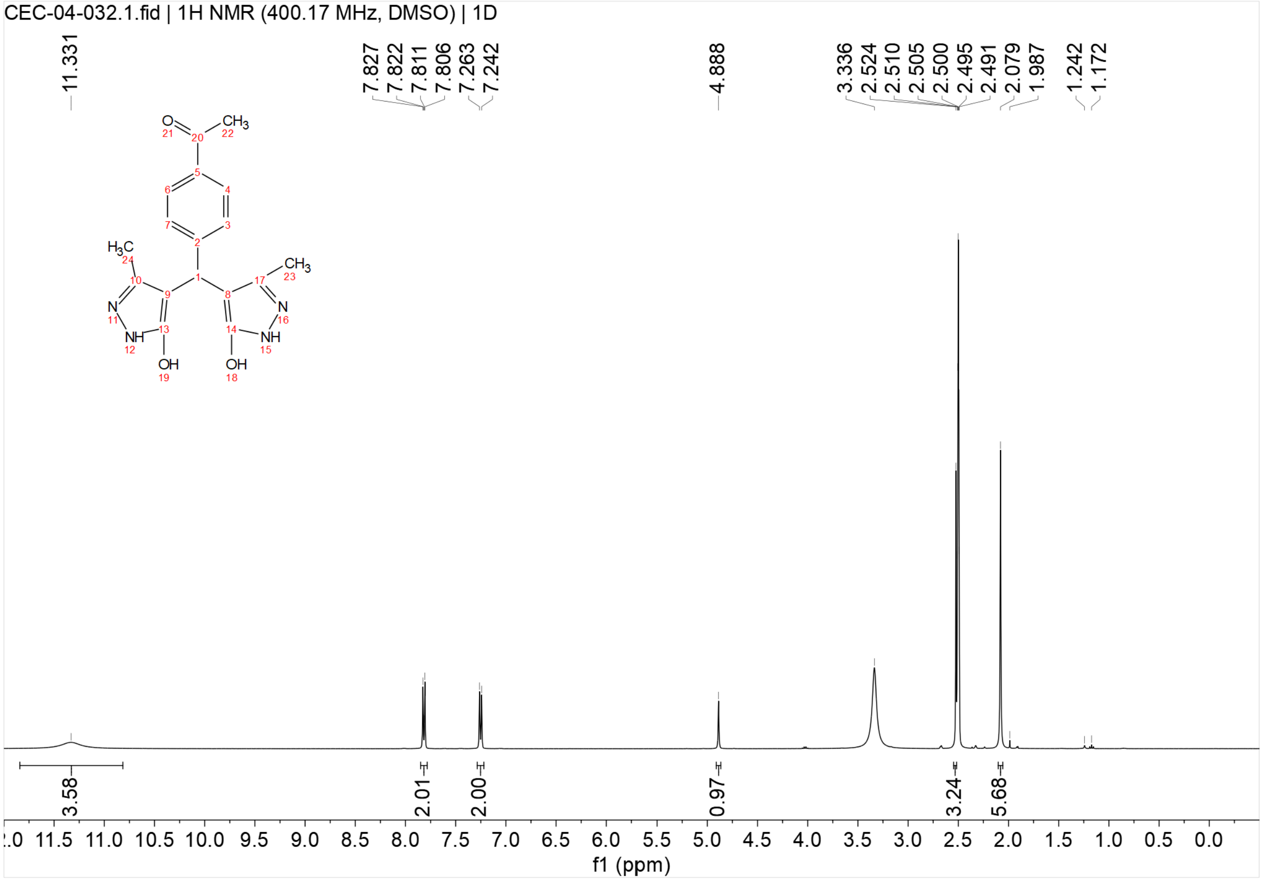


$^{13}C\{^{1}H\}$ NMR spectrum of **UP-7** (151 MHz, DMSO-$d_6$)

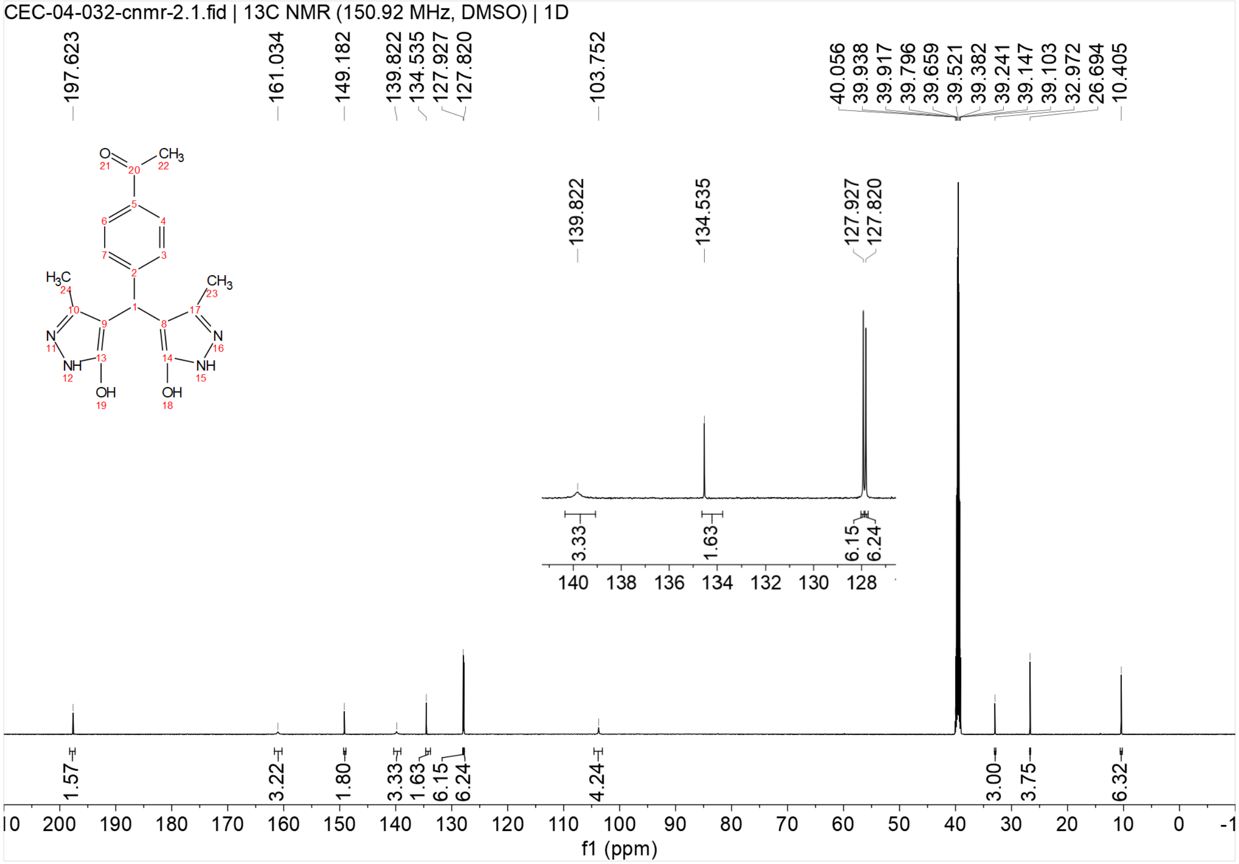


[1]H NMR spectrum of **UP-8** (600 MHz, DMSO-$d_6$)

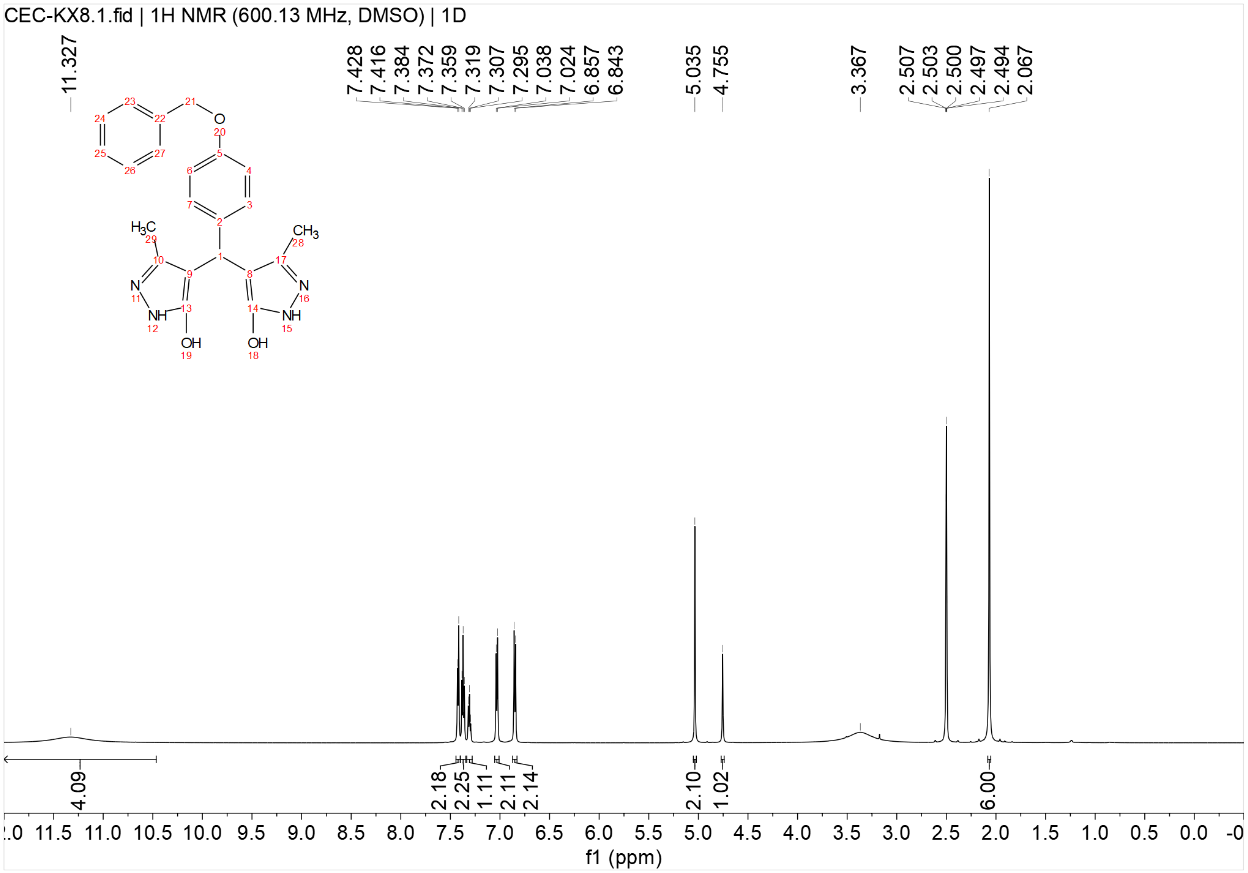


$^{13}C\{^{1}H\}$ NMR spectrum of **UP-8** (151 MHz, DMSO-$d_6$)

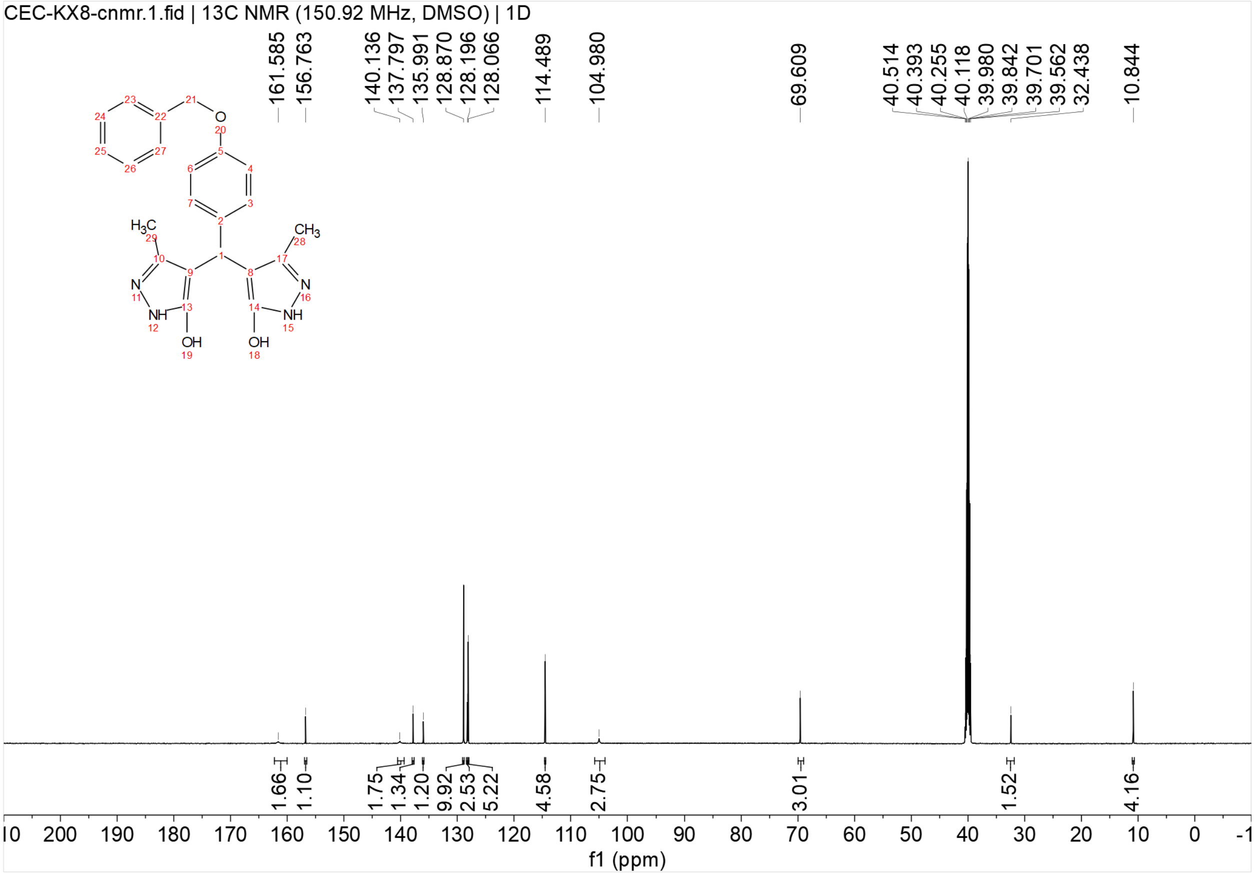

CEC-KX8-cnmr.1.fid | 13C NMR (150.92 MHz, DMSO) | 1D
161.585
156.763
140.136
137.797
135.991
128.870
128.196
128.066
114.489
104.980
69.609
40.514
40.393
40.255
40.118
39.980
39.842
39.701
39.562
32.438
10.844
1.66
1.10
1.75
1.34
1.20
9.92
2.53
5.22
4.58
2.75
3.01
1.52
4.16
f1 (ppm)

[1]H NMR spectrum of **UP-9** (400 MHz, DMSO-$d_6$)

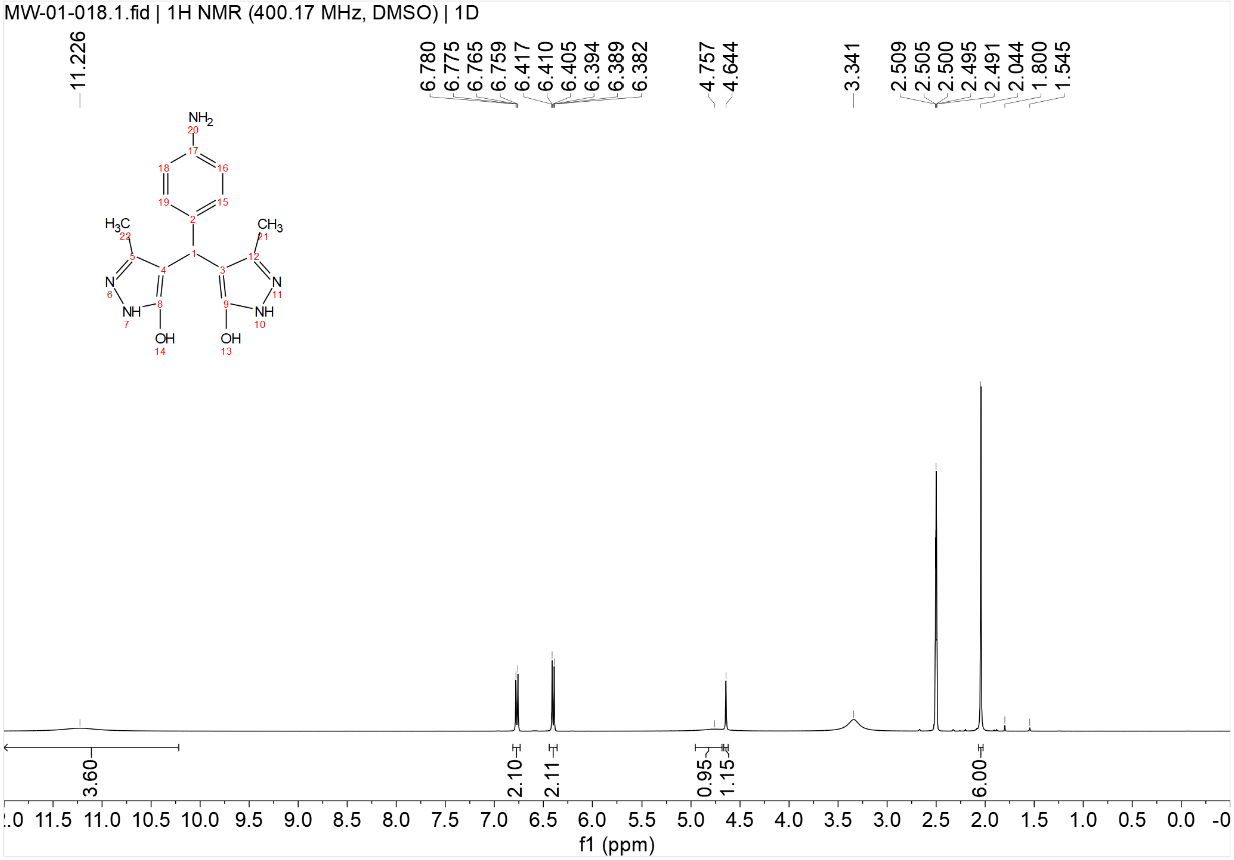


[13]C{[1]H} NMR spectrum of **UP-9** (151 MHz, DMSO-$d_6$)

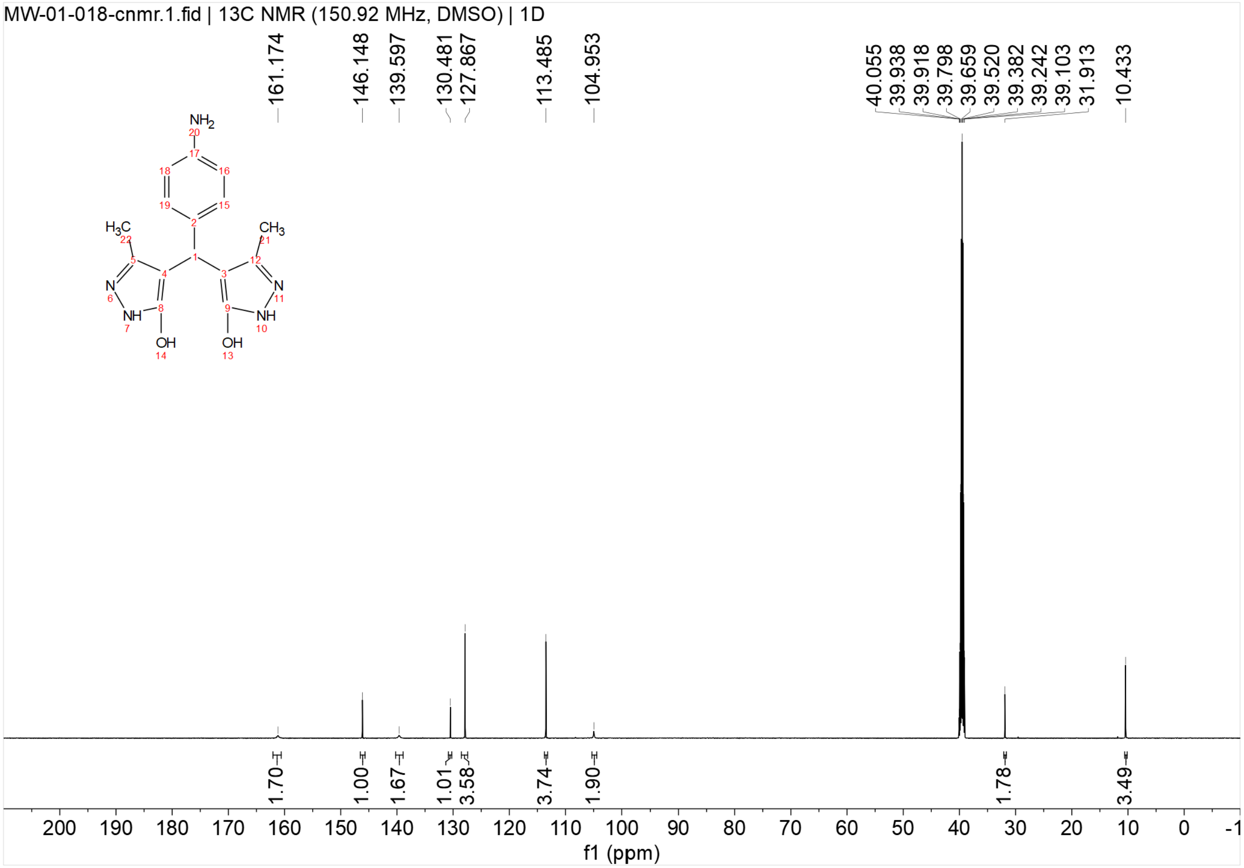


$^{1}H$ NMR spectrum of **UP-10** (600 MHz, DMSO-$d_6$)

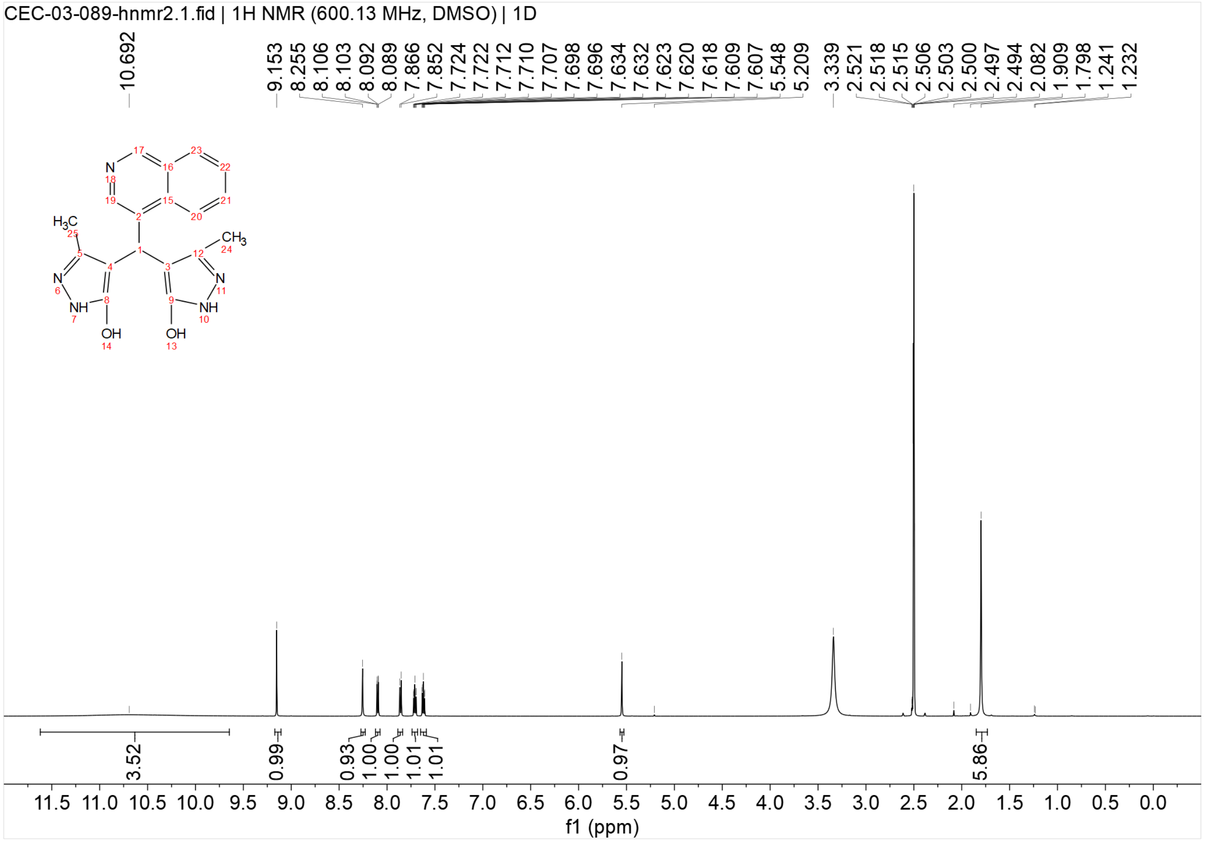

$^{13}C\{^{1}H\}$ NMR spectrum of **UP-10** (151 MHz, DMSO-$d_6$)

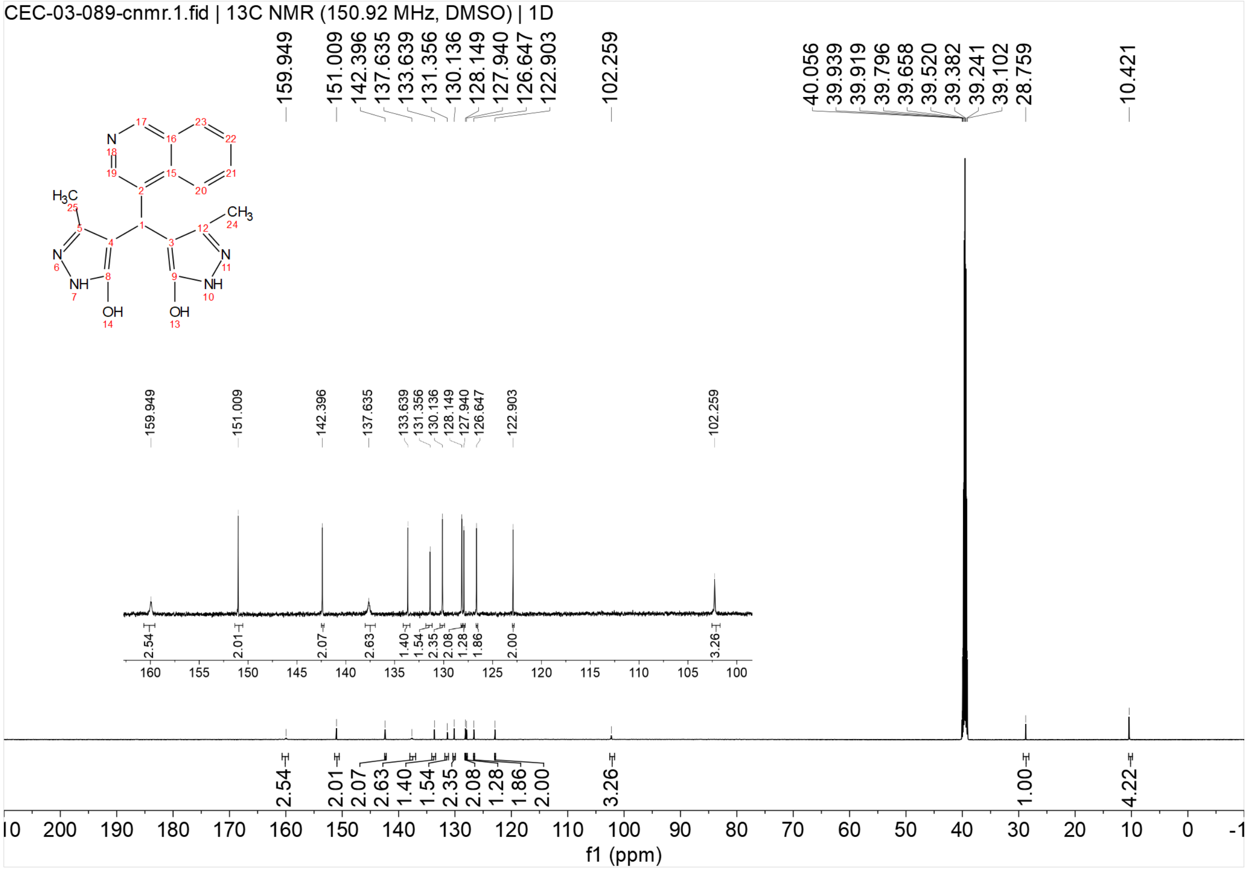

$^{1}H$ NMR spectrum of **UP-11** (400 MHz, DMSO-$d_6$)

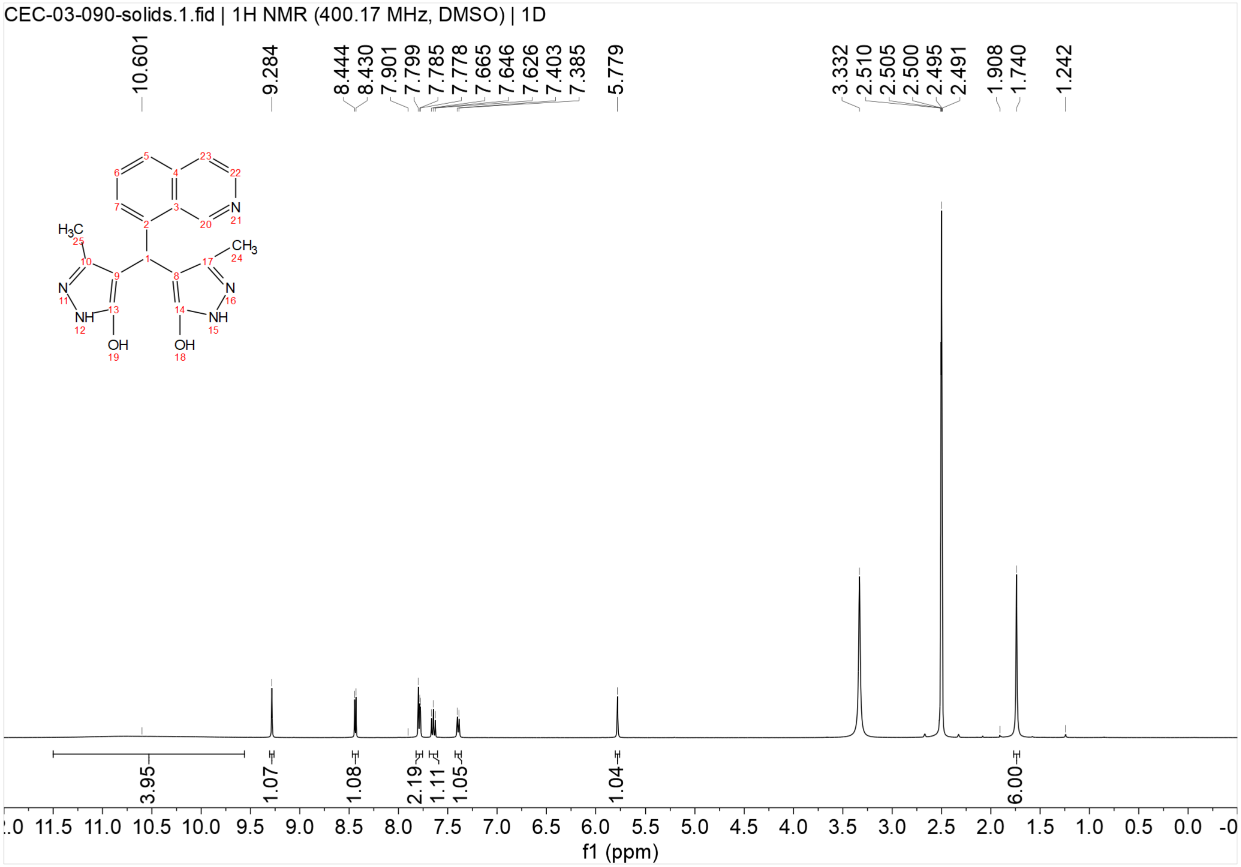


$^{13}C\{^{1}H\}$ NMR spectrum of **UP-11** (151 MHz, DMSO-$d_6$)

CEC-03-090-cnmr.1.fid | 13C NMR (150.92 MHz, DMSO) | 1D

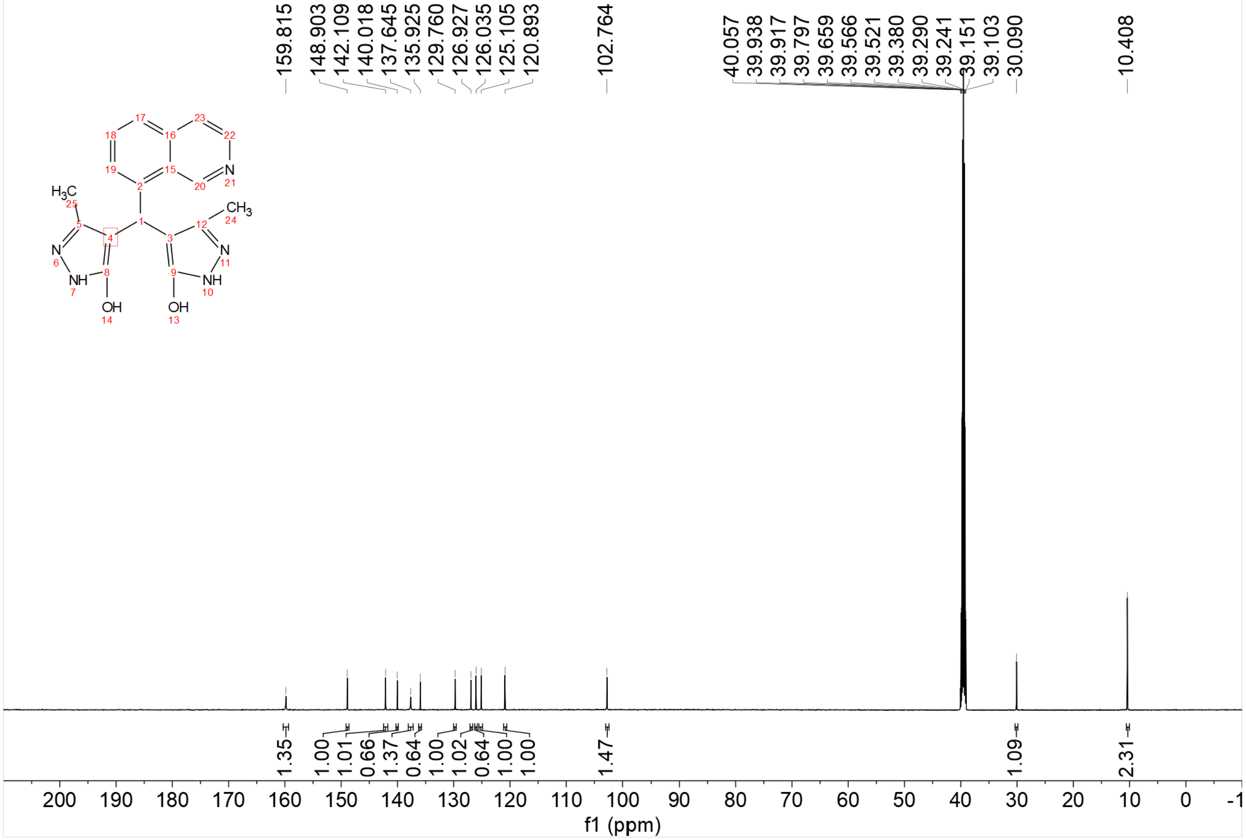

$^{1}H$ NMR spectrum of **UP-12** (400 MHz, DMSO-$d_6$)

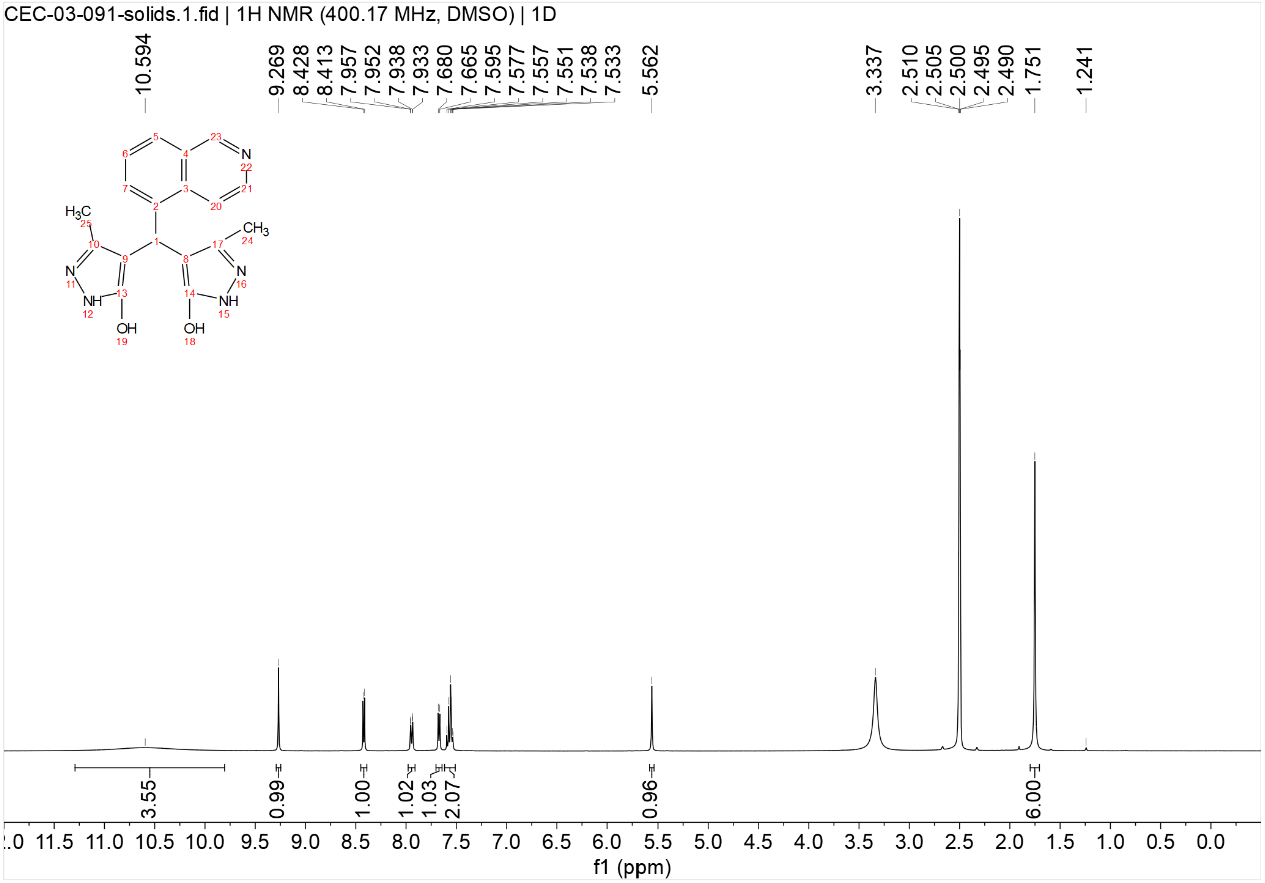


$^{13}C\{^{1}H\}$ NMR spectrum of **UP-12** (151 MHz, DMSO-$d_6$)

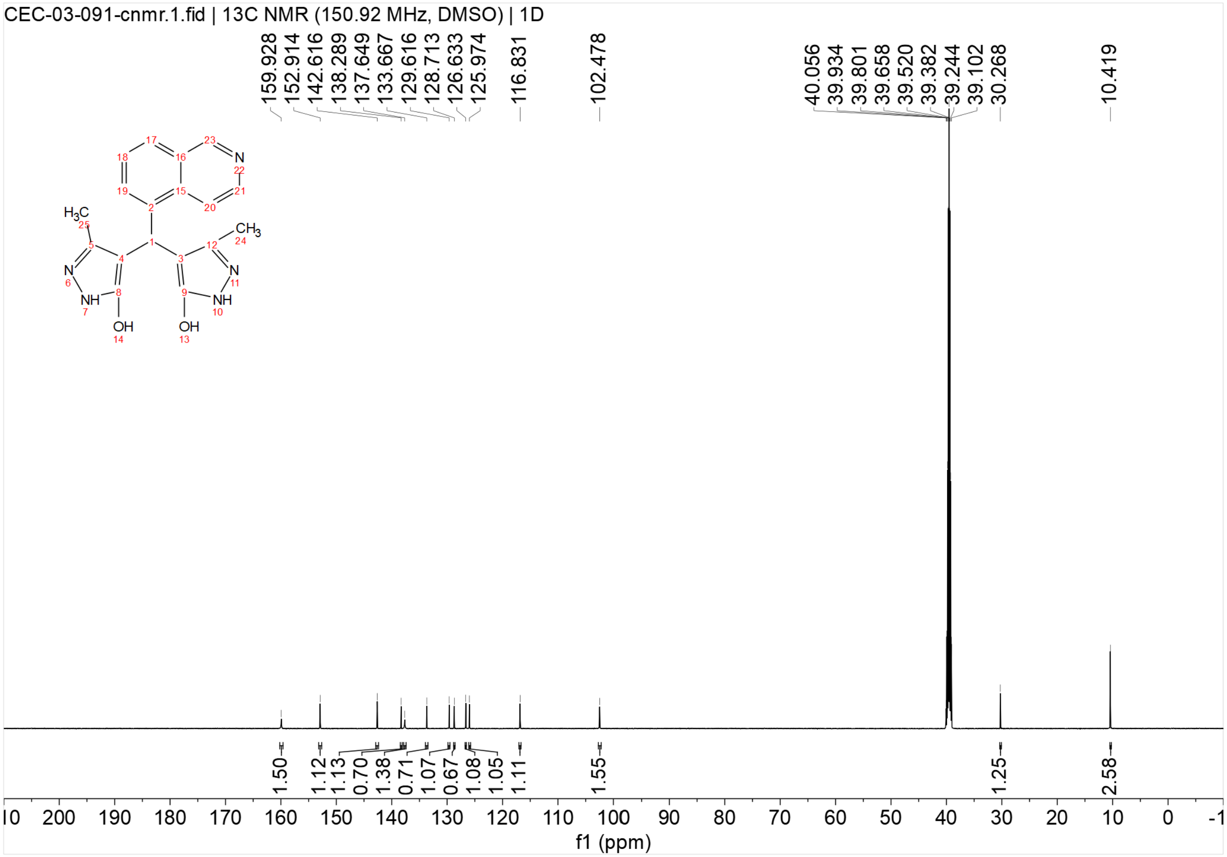


[1]H NMR spectrum of **UP-13** (400 MHz, DMSO-$d_6$)

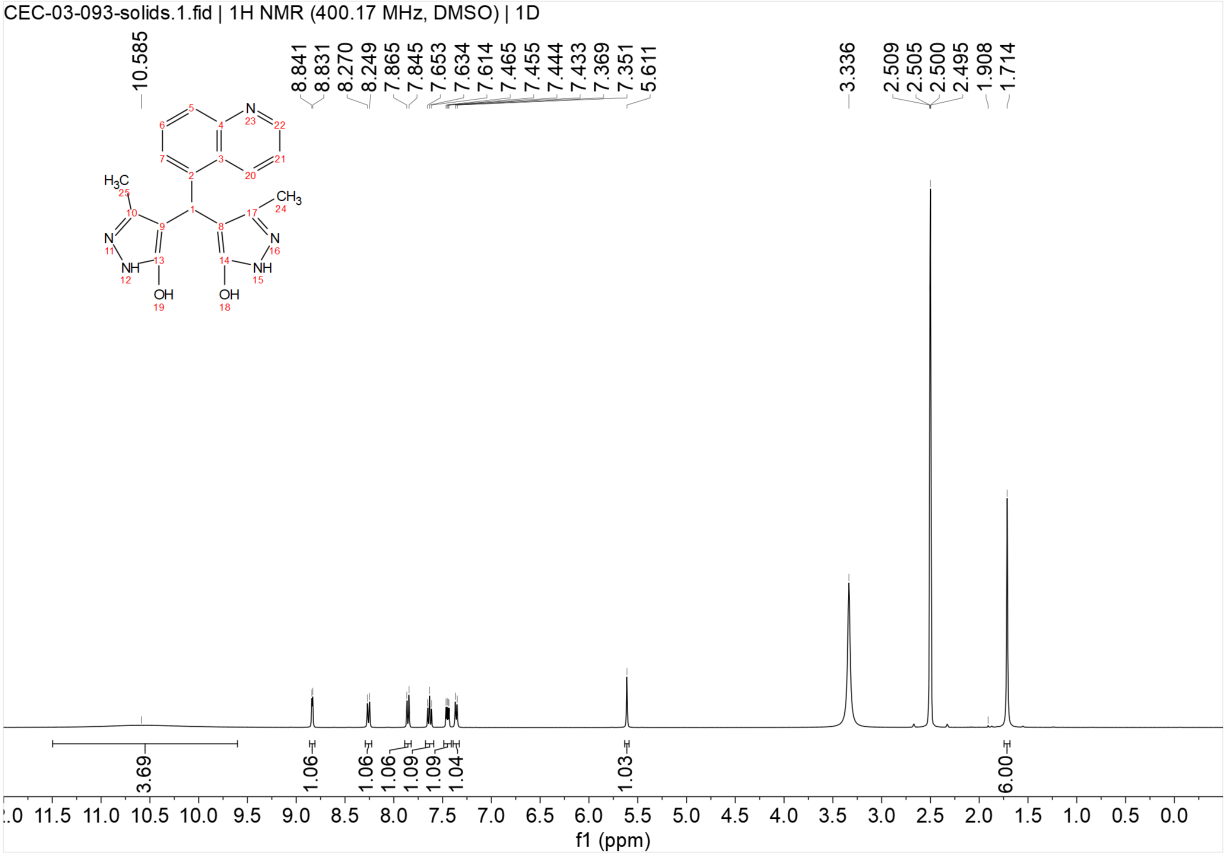

$^{13}C\{^{1}H\}$ NMR spectrum of **UP-13** (151 MHz, DMSO-$d_6$)

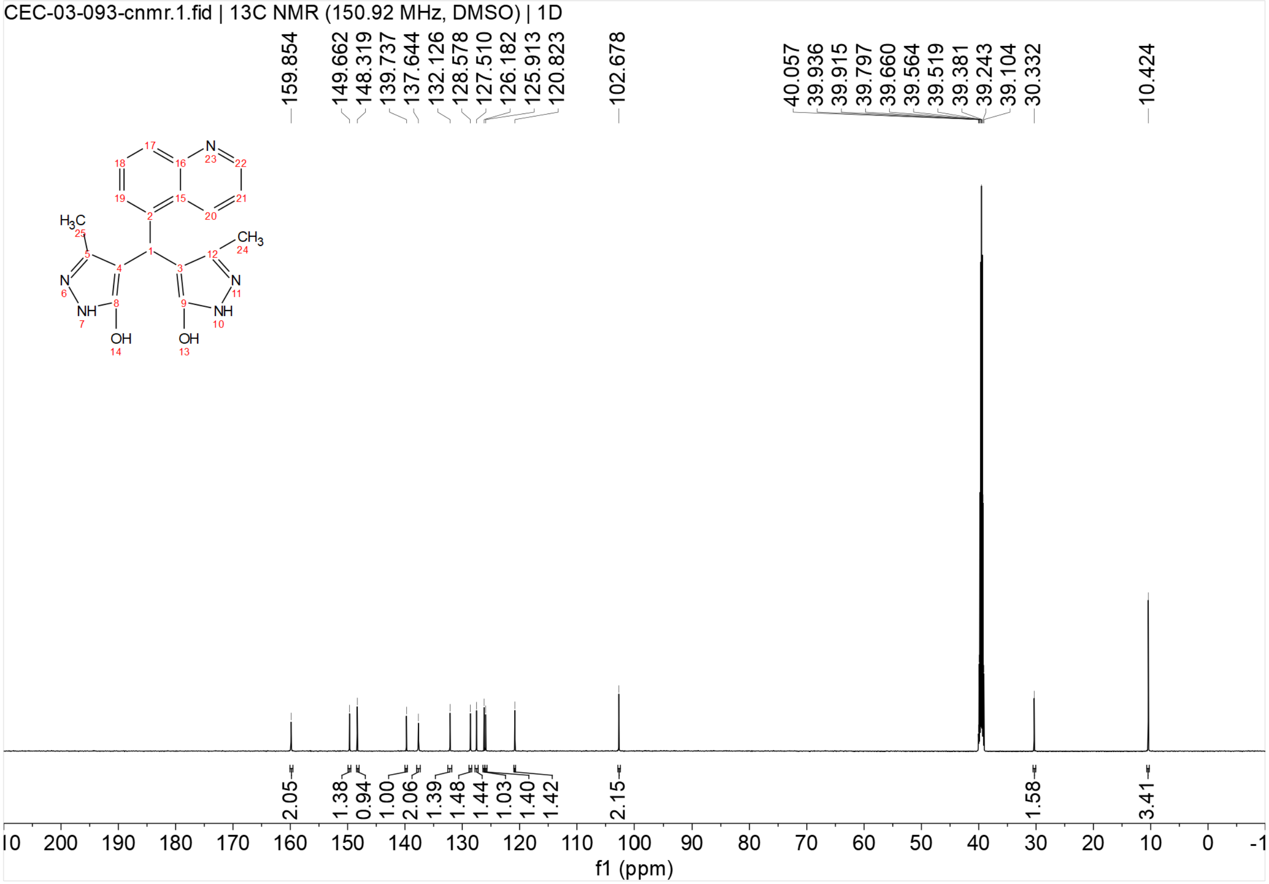

**Section E – Mass-spectrometry of lipid microbubbles loaded with compounds**

Mass spectrometry confirmation of C74 and UP-6 encapsulation in lipid microbubbles. Only C74 and UP-6 characteristic mass peaks were detected in microbubble samples.

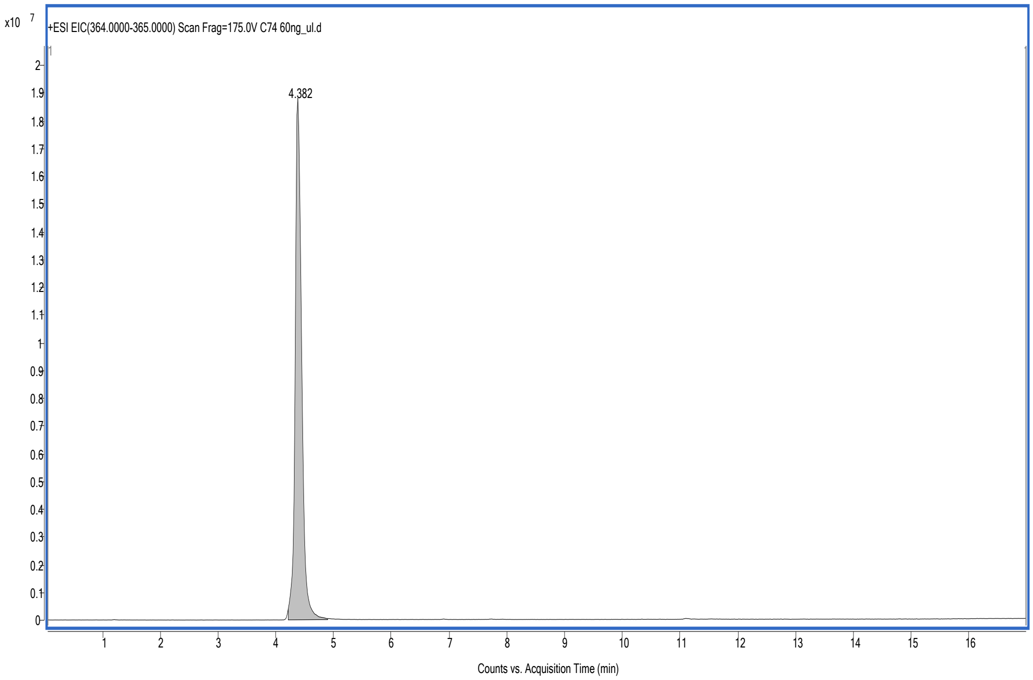


C74 Standard (Area = 148395494.17)

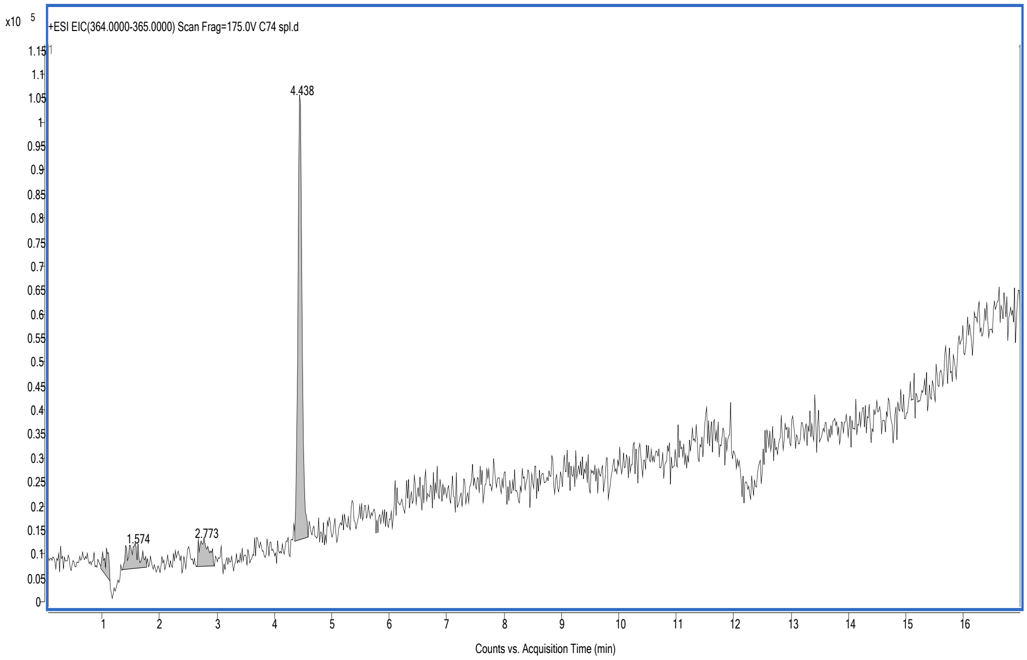


C74 Sample (Area = 470581.95)

Amount in Sample (ng/µL) = 0.190

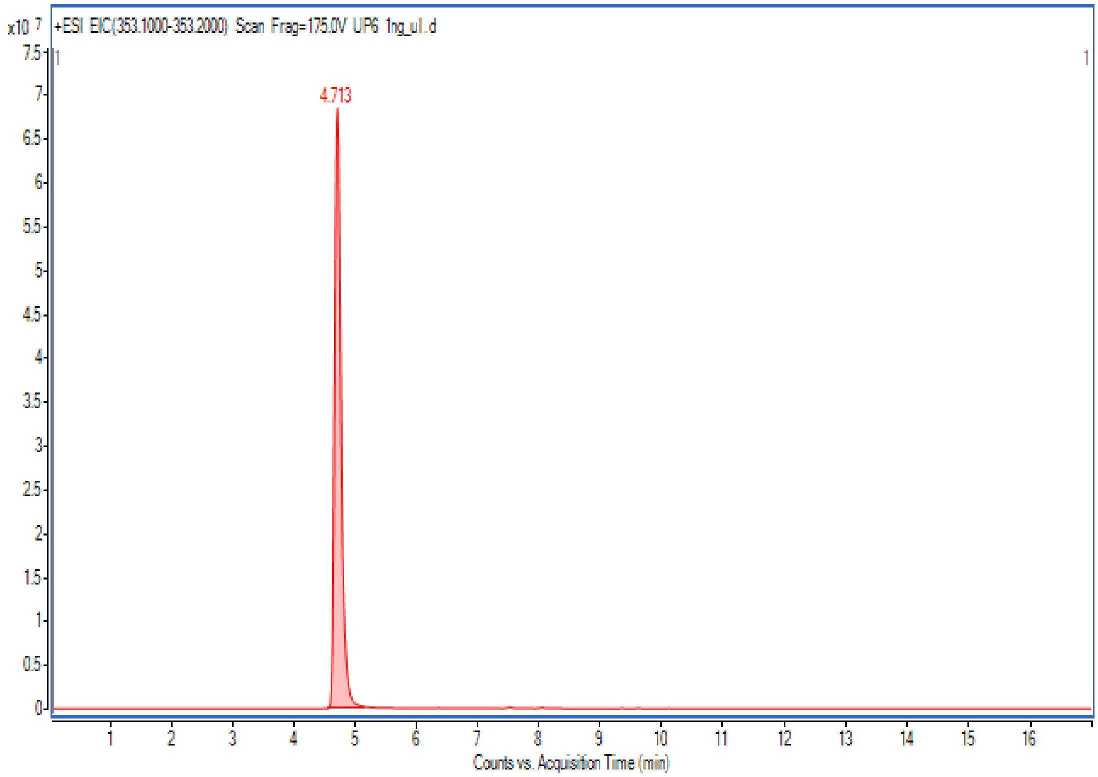


UP6 Standard (Area = 503056595)

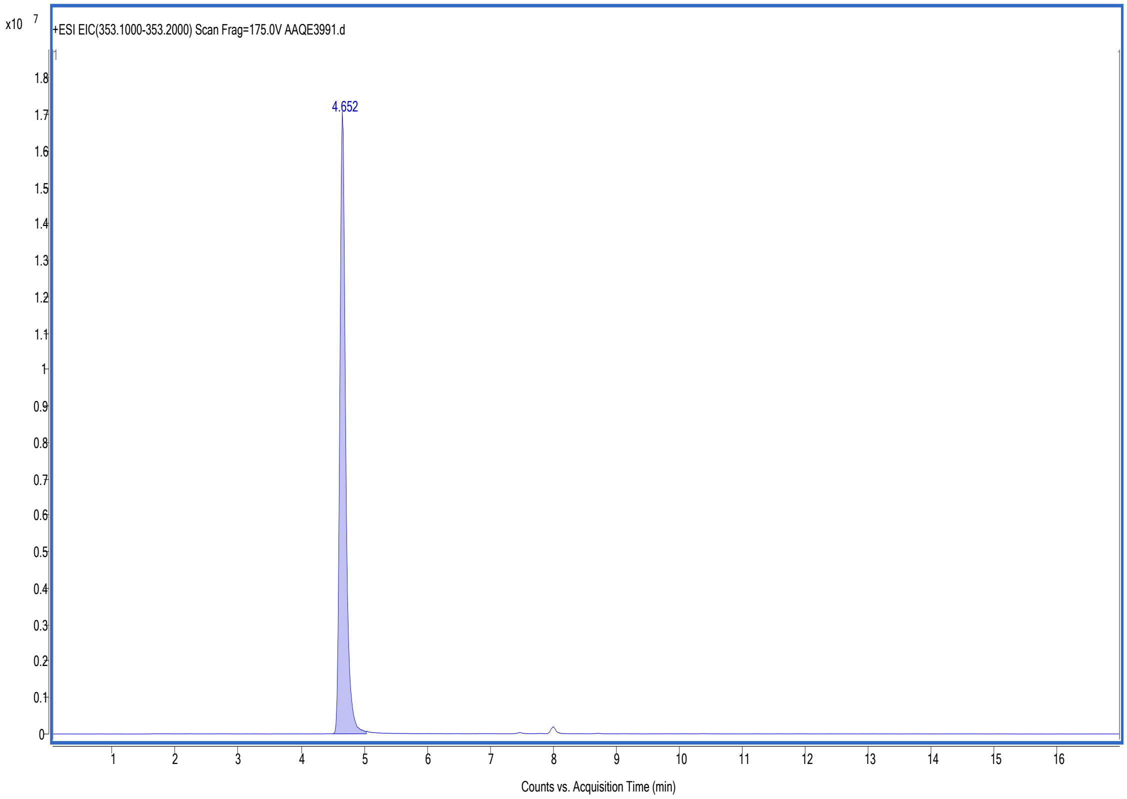


UP6 Sample (Area = 111178349)

Amount in Sample (ng/µL) = 0.221
